# Mapping Senescence Landscapes in Metabolic Liver Disease Progression

**DOI:** 10.64898/2026.11.12.766040

**Authors:** Elizabeth L. Schmidt, Lee Williams, Samuel T. Peters, Rahagir Salekeen, Natalia Calixto Mancipe, Linshan Laux, Mickayla DuFresne-To, Christine Henzler, Sayeed Ikramuddin, Jinhua Wang, Tim Meyer, Steven Johnson, Constantin Aliferis, Byoung Uk Park, Oyedele Adeyi, Erich Kummerfeld, Paul D. Robbins, Laura J. Niedernhofer, Andrew C. Nelson

## Abstract

Metabolic dysfunction-associated steatotic liver disease (MASLD) affects more than 30% of the population, often progressing from reversible steatosis to irreversible fibrosis and cirrhosis. Although cellular senescence is implicated in driving MASLD, the evidence in human tissue and the mechanism are lacking. Here, we apply spatial transcriptomics and scRNAseq to human liver samples spanning healthy to fibrotic disease to address these knowledge gaps. Using a combination of cell stress, cell cycle arrest, extracellular matrix (ECM) remodeling, and inflammatory genes, senescent microenvironments termed “SenSpots” were identified by Visium transcriptomics that predominantly localized to portal tracts and fibrotic septa across all specimens. Functional profiling revealed a fundamental phenotypic divergence: SenSpots in healthy liver displayed a physiological wound-healing and tissue-development program, whereas SenSpots in fibrotic tissue expressed genes related to inflammation, stress, and ECM-disassembly. Spatial deconvolution and network expansion analyses demonstrated that SenSpots act as epicenters for TGF-*β*, NOTCH, complement, and ECM-reorganization signaling, which spreads into the surrounding tissue microenvironment as disease advances. Furthermore, scRNAseq data validated the Visium results and identified a previously unappreciated population of senescent Schwann cells in the diseased liver niche. These findings map the spatial origination and paracrine propagation of pathological senescence and identify senotypes associated with liver fibrosis.

## Introduction

Chronic liver disease is a major cause of global morbidity and mortality, driven predominantly by metabolic dysfunction–associated steatotic liver disease (MASLD) and its progressive inflammatory phenotype, metabolic dysfunction–associated steatohepatitis (MASH). While many individuals with hepatic steatosis remain clinically stable, a substantial subset progresses to severe fibrosis, cirrhosis, and hepatocellular carcinoma. Cirrhosis reflects an advanced stage marked by architectural collapse, portal hypertension, and liver failure, for which curative options beyond liver transplantation remain severely limited.^1,2^ This is particularly critical in aging contexts, where limited donor availability and compromised viability drive disproportionate morbidity and mortality in patients over 65.^3,4^ Although genetic, metabolic, and inflammatory drivers of liver disease have been extensively profiled,^2,5–7^ the work is largely correlative rather than mechanistic.

Cellular senescence has emerged as a central driver of chronic liver injury, directly linking metabolic stress, aging and inflammation to fibrotic remodeling^8–12^. Senescent cells undergo stable cell-cycle arrest paired with anti-apoptotic survival signaling and extensive secretory reprogramming, collectively termed the senescence-associated secretory phenotype (SASP). In human liver disease, senescent cell traits have been described across diverse cell types. However, existing studies relied heavily on bulk transcriptomics or dissociative single-cell RNA sequencing (scRNA-seq).^9,13,14^ Because enzymatic dissociation strips cells of their histological context, these approaches preclude direct interrogation of how senescent cells are organized within anatomical hepatic zones, how they interface with fibrotic septa, and how their paracrine signaling contributes to disease progression. Furthermore, hepatocyte-centric studies obscure critical non-parenchymal senescence, ignoring recent progress in establishing senescence-like phenotypes in Kupffer cells and liver endothelium.^7,15^

The liver exhibits a highly ordered spatial architecture, organized along a porto-central metabolic zonation axis that dictates oxygen gradients, nutrient flux, regeneration and regional susceptibility to injury.^16^ Dissecting how senescence disrupts this architecture requires platforms that capture whole-transcriptome signaling while preserving histological and anatomical context. Spatial transcriptomic approaches overcome key limitations of prior methodologies by enabling transcriptome-wide spatial profiling directly from formalin-fixed, paraffin-embedded (FFPE) biopsies. This allows systematic integration of molecular transcriptomics with histopathological landmarks across the human liver disease spectrum.

Here, we leveraged Visium CytAssist to map senescent cells in healthy and diseased human liver. By applying a multi-category senescence signature, we identified senescence microenvironments (“SenSpots”), which localized primarily to portal tracts and fibrotic niches across all tissue stages. Functional profiling revealed a clear distinction in senescence phenotypes: SenSpots in healthy liver displayed a wound healing and tissue development program, while those in fibrotic tissue displayed a pro-inflammatory and ECM-disassembly phenotype. scRNA-seq confirmed the spatial results and identified specific cell populations, including senescent hepatic stellate cells and myofibroblasts as key drivers, while also identifying a novel population of senescent Schwann cells in liver. Combining the datasets revealed paracrine signaling emanating from SenSpots, spreading senescence features likely via intercellular TGF-β, NOTCH, ECM remodeling, and complement signaling pathways.

## Results

### Cohort Description

We examined formalin fixed paraffin embedded (FFPE) human liver biopsies using Visium CytAssist spatial transcriptomics (10x Genomics). This platform captures a uniform grid of spatially-resolved micro-regions (55 µm diameters) containing on average 5-30 cells (**Figure 1A**). Eight liver biopsies (2 male, 6 female) were processed with Visium CytAssist from subjects ranging in age from 32-50 yo with a spectrum of liver disease defined by standard pathologic grading (**Figure 1B**). There were 4 healthy tissue samples (normal, age range 32-49 yo), one with mild lobular inflammation, one with mild steatosis, one with a combination of mild steatosis, inflammation, and fibrosis, and one cirrhotic sample (fibrosis score = 4). The raw probe sequencing reads were spatially mapped using the Space Ranger pipeline (**Supplemental Table 1**), and artifact removal was performed using BLADE.^1^

**Figure 1.**
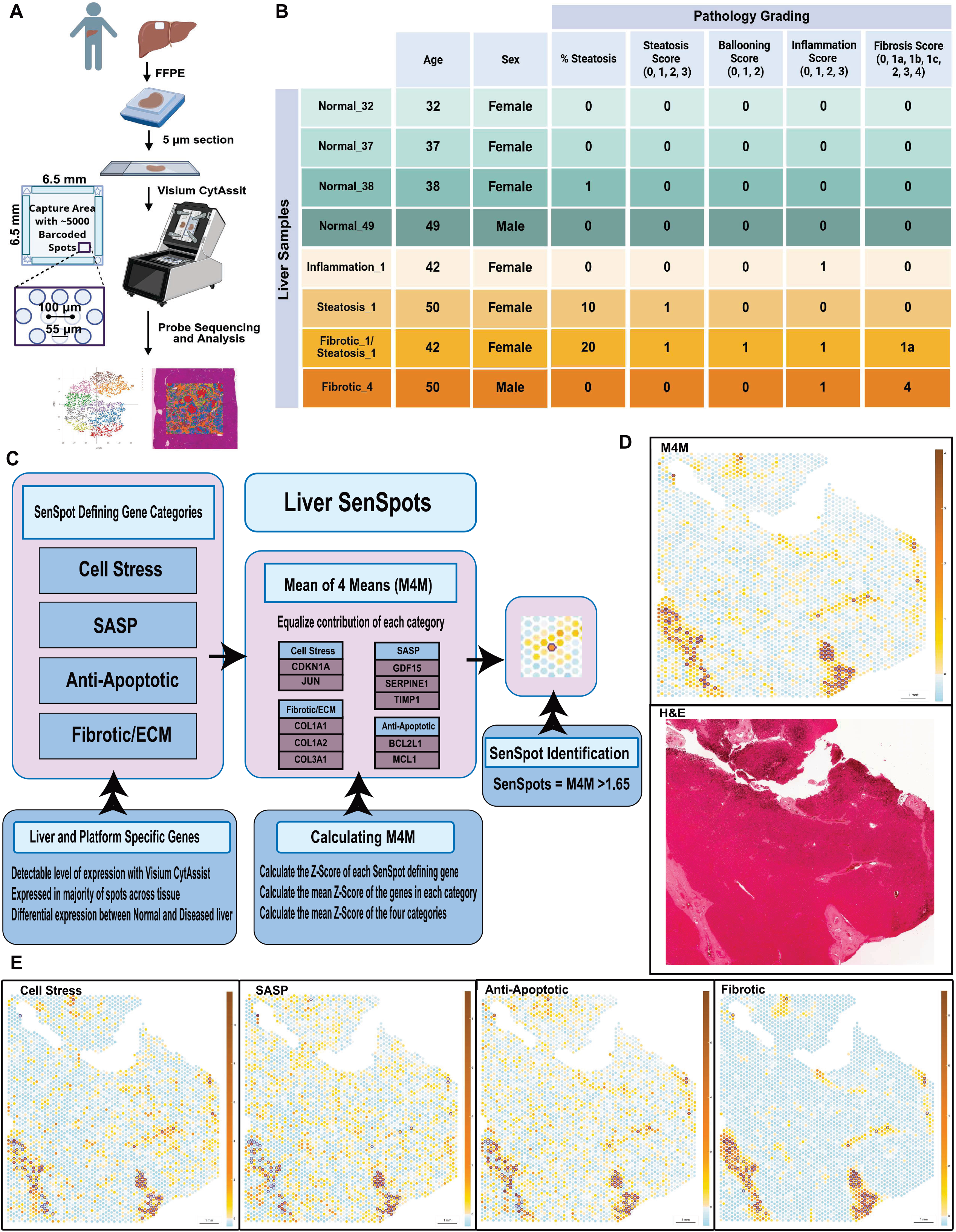
Liver SenSpot Identification. **A**. Human liver biopsies were formalin fixed and paraffin embedded. 5***μ***m sections were processed with Visium CytAssist. The Visium 6.5 x 6.5 mm capture area contains >5,000 spots of 55 µm diameter with 100 µm distance from center to center of adjacent spots. The CytAssist instrument transfers the transcriptomic probes from the histology slide to the Visium capture slide, retaining the spatial location of transcripts **B.** Pathology grading of liver biopsies from the 8 subjects **C.** SenSpots were defined by increased expression of 10 genes from 4 senescence-related categories. The mean of 4 means (M4M) was determined using the mean z-score of each senescence category and then calculating the mean z-score of all 4 categories ≥1.65. **D.** Image of Visium spot heatmap of the M4M for Inflammation_1 liver sample (top) and H&E (bottom) **E.** Spatial heatmaps of the mean z-score for each of the 4 senescence gene categories.

### SenSpot Identification and Clustering

Our goal was to establish a compact, cross-pathology marker panel that captured the multi-faceted nature of senescent cell features to consistently identify tissue-level ‘SenSpots’ across heterogeneous liver samples. Senescence-associated gene expression from four biologic senescence-associated processes were interrogated: cell stress, senescence associated secretory phenotypes (SASP), anti-apoptosis, and extracellular matrix (ECM) remodeling/fibrosis to identify senescence niches within liver tissue (**Figure 1C**). Cell stress, included genes associated with cell cycle arrest/control, the DNA damage response, and chromatin structure. The SASP category included cytokines, chemokines and growth factors commonly found secreted as part of the SASP. The anti-apoptotic category included BCL2 family genes often upregulated in senescence to maintain cell survival. The ECM/fibrotic category identified features of a remodeling tissue.

To identify genes that most reliably detected senescence in human liver, we compared the variance in gene expression across the tissue for the four categories using healthy and cirrhotic liver samples (**Supplement Table 2**). Genes were selected based on three criteria: (1) reliable detection on the Visium CytAssist platform, (2) broad baseline expression across the majority of spatial spots in diseased tissue, and (3) high expression variance between normal and diseased samples. Using these criteria, we identified the 10 best-performing SenSpot genes representing each of the four senescence categories, which were: *CDKN1A* and *JUN* (cell stress); *GDF15*, *SERPINE1*, and *TIMP1* (SASP); *BCL2L1*, *MCL1* (anti-apoptotic); and *COL1A1*, *COL1A2* and *COL3A1* (ECM). To ensure contribution of genes from each category, we utilized the mean z-score of all 4 categories, or the mean of 4 means (M4M) calculation with a threshold of >1.65 to identify SenSpots (**Figure 1C**). M4M for SenSpot identification was based on individual sample normalization, making the threshold determination specific for each sample.

Utilizing this method, we identified liver specific SenSpots within each tissue sample. A spatial heatmap from sample Inflammation_1 of M4M genes with SenSpots illustrated their localization within the fibrous tissue of portal tracts apparent on the Visium H&E image **(Figure 1D**). Unique, non-overlapping gene expression patterns were observed in the spatial heatmaps for the 4 distinct gene categories (**Figure 1E**). However, there was also a strong overlap of SenSpots defined by each category. The pattern of SenSpot concentration within portal tracts and areas of fibrosis was observed in both in normal and diseased tissues across the cohort (**Extended Data Figures 1-2**).

To further test whether any of the four individual senescent cell hallmark signatures sufficiently identified SenSpots, the expression of each gene category was analyzed separately (**Extended Data Figure 3A**). Candidate niches identified by only single-category gene enrichment showed inconsistent gene expression in the other three hallmarks, in contrast to SenSpots identified by the four-category M4M model which exhibited robust upregulation across all ten target genes. Additionally, SenSpots identified with M4M demonstrated upregulation of an expanded group of established senescence markers beyond the M4M gene set (**Extended Data Figure 3B**). Importantly, the correlation of M4M gene expression with each other senescence-associated genes strengthened progressively with disease severity, underscoring the link between senescence activation and disease progression (**Extended Data Figure 3C**). In summary, we developed a multi-category senescence signature (M4M) to identify SenSpots across heterogeneous tissue contexts. By benchmarking our 10-gene panel against single-category subsets, we demonstrated that integrating all four functional categories is advantageous for capturing senescent tissue phenotypes. Furthermore, the correlation of M4M gene expression scaled with disease, establishing this gene list as a transcriptomic indicator of liver pathology.

### Characterization of SenSpots

We next identified differentially expressed genes (DEGs) upregulated in SenSpots compared to non-senescent tissue regions in each individual sample. Subsequently, genes upregulated within SenSpots from healthy tissue samples (n=4 donors) were cross-referenced to those upregulated in SenSpots of the cirrhotic liver sample (Fibrotic_4) to identify unique and shared SenSpot-associated genes (**Figure 2A**). Forty-four DEGs were upregulated only in the Fibrotic_4 tissue and led to enriched pathways associated with pro-inflammatory immune response and ECM disassembly. Further mapping and Markov clustering of the DEG interactome suggested two separate clusters of genes enrich these two processes, namely centered on immune regulation (*LAPTM5*, *LYZ*, *CD163*, *LCP1*, *AIF1*) and ECM remodeling (*MMP*s and *ITGB*s). Ninety-two DEGs were upregulated only in the normal samples and were enriched for ECM organization, blood vessel morphogenesis, cell differentiation, and wound healing. Forty-five DEGs common to SenSpots from both healthy and cirrhotic tissue were associated with collagen fibril assembly, regulation of cell migration, and calcium-dependent protein binding pathways, with interactomes centered on various collagen and other ECM gene products.

**Figure 2.**
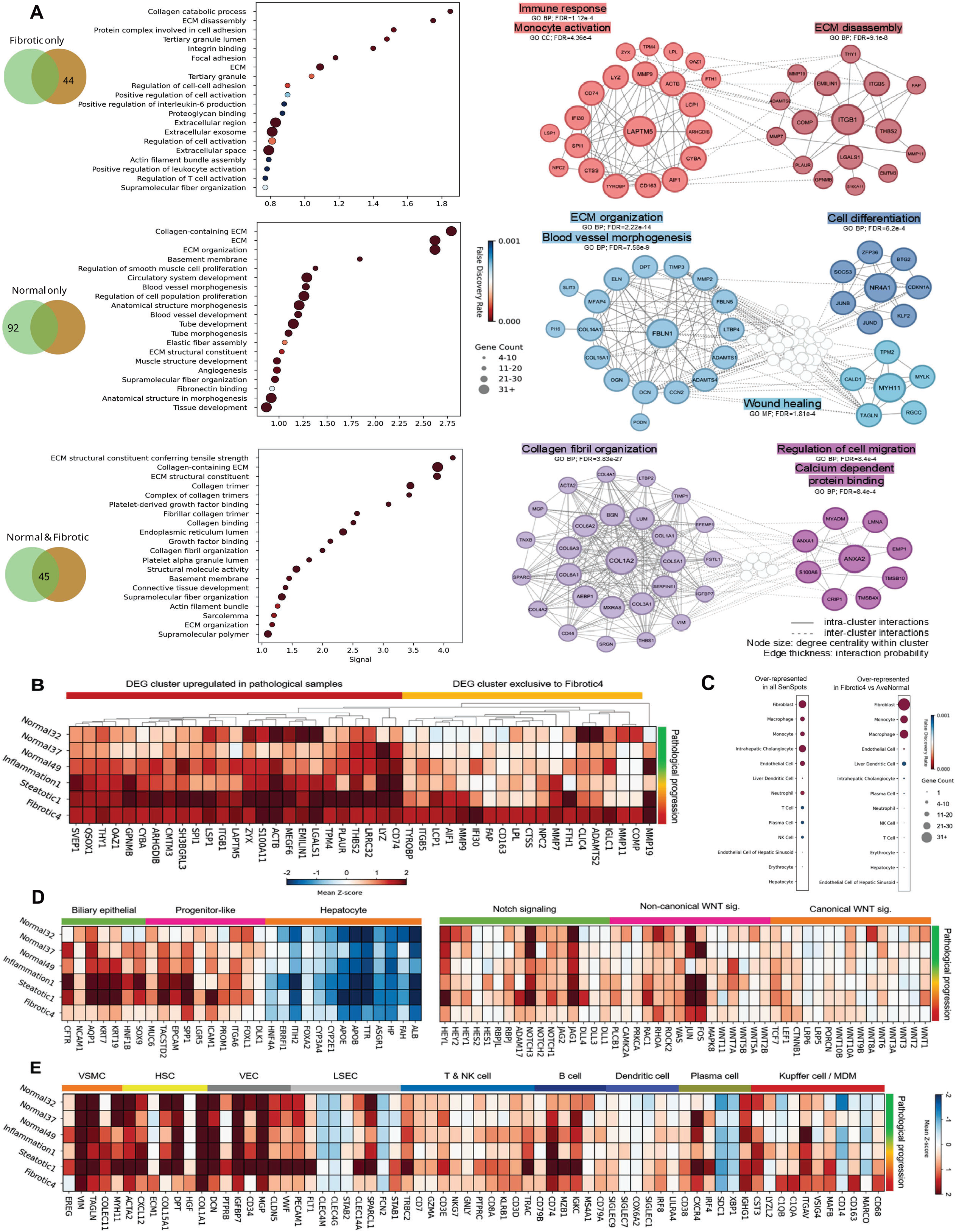
Visium SenSpot characterization. **A.** Bubble chart of enriched GO Biological Process (GO:BP) and Molecular Function (GO:MF) terms. Top 20 GO enriched pathways are visualized from upregulated DEGs in SenSpots shared between Ave_Normal and Fibrotic_4, unique to Ave_Normal, and unique to Fibrotic_4. X-axis: enrichment signal; color: False Discovery Rate (FDR); bubble size: observed gene count bins. Major Markov clusters (MCL) of protein-protein interaction network of exclusive differentially expressed gene (DEG) nodes and their physical/functional associations (edges) from SenSpot genes overlapping for Normal & Fibrotic, unique to Normal, and unique to Fibrotic visualized as clustered radial diagrams. Edge width represents interaction confidence score and node size represents degree centrality within network. **B.** Heatmap shows mean empirical Z-scores across all samples for DEGs up in Fibrotic_4 after filtering for Ave_Normal. Gene columns are clustered by expression similarity **C.** Enrichment for the *Tabula Sapiens* dataset for cell type signature deconvolution in all SenSpots (mean Z-score>=1.5) and Fibrotic_4 vs Ave_Normal (mean Z-score >=1.5 in Fibrotic_4 and <1.5 in Ave_Normal). **D-E.** Heatmaps displaying SenSpot mean Z-scores for genes across four different liver samples with increasing pathology from top to bottom based on the color-coded annotation bar (right). **D.** Genes associated with hepatic differentiation from bipotent progenitors to mature hepatocytes or cholangiocytes, and by Notch or WNT signaling (canonical and non-canonical) with rows grouped by the color-coded annotation bar (top). **E.** Genes associated with cell identity by the color-coded annotation bar (top). Monocyte-derived macrophage (MDM), Natural killer (NK), liver sinusoidal endothelial cells (LSECs), hepatic stellate cells (HSCs), vascular smooth muscle cells (VSMCs).

Given this result of different senotypes between normal and cirrhotic tissue SenSpots, we explored the dynamics of these DEGs across additional samples representing histologic disease progression. Two major clusters of DEGs were identified: (i) genes that scaled gradually with pathological progression and (ii) genes that were exclusive to cirrhotic SenSpots (**Figure 2B**). The first cluster included senescent cell surface markers (*GPNMB*, *PLAUR*), functional drivers of an activated mesenchymal phenotype expressed by cancer-associated fibroblasts (CAFs) and mesenchymal stem/stromal cells (*THY1*, *THBS2*), TGF-β signaling regulator (*LRRC32*), and scar-associated macrophage activators (*SPI1*, *CD74*, *LYZ*). The second cluster of genes in cirrhotic SenSpots included robust upregulation of matrix-degrading metalloproteinases and remodeling enzymes (*MMP*s, *ADAMTS2*, *FAP*) and markers of disease-associated macrophages and immunoglobulins (*CD163*, *TYROBP*, *AIF1*, *IGLC1*), suggesting local ECM disassembly and chronic inflammation.

Overrepresentation analysis of cell-type-specific signatures in SenSpots further corroborated their localization predominantly in non-parenchymal portal tracts and/or fibrotic regions. SenSpot DEGs across all samples showed strong enrichment for fibroblasts, myeloid cells, intrahepatic cholangiocytes, endothelial cells and neutrophils, while hepatocyte- and sinusoidal endothelium-specific enrichment was low (**Figure 2C**). Fibroblast and myeloid cell-specific signatures were further upregulated in SenSpots from cirrhotic vs normal samples. Collectively, spatial profiling across liver disease states suggested that SenSpots arise in non-parenchymal niches driven by matrix remodeling and homeostatic processes, but their downstream signaling may diverge according to disease stage contexts (**Figure 2A-C**). While normal tissue SenSpots exhibited a phenotype enriched in genes related to tissue development and wound-healing, cirrhotic SenSpots were characterized by pro-inflammatory stress responses and ECM breakdown.

Hepatic homeostasis and resolution of transient stress signaling depends on hepatocyte regeneration and function. Age-related loss of regenerative capacity enables non-parenchymal cell types to infiltrate the normal tissue architecture. Given the lack of SenSpot enrichment in hepatocytes (**Figure 2C)**, an extended list of genes associated with hepatobiliary regeneration transdifferentiation suggested a bias towards progenitor like/biliary cholangiocyte markers in SenSpots, particularly in inflamed /steatotic tissues but dampened in the end-stage fibrotic sample (**Figure 2D**). A few possible scenarios emerge from these signatures: (i) senescent cell features are primarily expressed in non-parenchymal cells and hepatocytes are excluded from these niches, (ii) hepatocyte regeneration is impaired in SenSpots leading to elimination of hepatocytes, or (iii) hepatocyte transcriptional plasticity may lead to loss of functional identity in SenSpots. While transcriptional snapshots alone cannot resolve these non-exclusive scenarios, drivers of alternative regenerative processes such as JAG-NOTCH, and non-canonical WNT signaling pathways were preferentially upregulated in SenSpots. However, hepatic and biliary regeneration were dampened in late-stage fibrosis/cirrhosis. SenSpots in all samples were enriched for vascular endothelial cells (VEC), B cells, and hepatic stellate cells (HSCs), while liver sinusoidal endothelial cell (LSEC) markers were downregulated (**Figure 2E**). Myeloid cells and dendritic cell markers were unique to steatotic and fibrotic SenSpots. Thus, SenSpots showed marked loss of parenchymal and sinusoidal endothelium enrichment and, in late-stage liver disease, signaling from pro-inflammatory myeloid cells indicative of stromal activation and ECM disassembly consistent with loss of hepatic regenerative capacity in end-stage cirrhosis

### SenSpot Localization and Zonation

Localization of the SenSpots can determine the effects they might have on liver function and regeneration. The liver lobule is a repeating structural unit centered on a central vein, with portal tracts containing the portal vein, hepatic artery, and bile duct located at each corner (**Extended Data Figure 4A**). Blood flows from the portal tracts through hepatic sinusoids toward the central vein, creating metabolic zonation (periportal zone 1, midlobular zone 2, and pericentral zone 3). This flow establishes an oxygen gradient that is highest near the portal tracts and progressively decreases toward the central vein.^17,18^ The SenSpots in the M4M spatial map of the Infammation_1 liver sample were clustered in portal tracts with collagen deposition represented by blue staining in the trichrome image (**Extended Data Figure 4B**). Magnification of a SenSpot cluster area highlighted a portal tract and bile duct with dense collagen fiber deposition in the tract itself, but no collagen extension into the hepatic interface and absence of substantial portal inflammation. Zone-specific gene expression was used to specify hepatocyte pericentral and periportal regions of the tissues (**Extended Data Figure 4C**). The majority of SenSpots occurred in areas with downregulation of functional markers for both periportal and pericentral cell identity. Pathologist annotation of SenSpot localization confirmed they were located predominantly in portal tracts (**Extended Data Figure 4D-E**), where niches are dominated by biliary cholangiocytes, stromal cells and progenitor-like hepatic cells. All SenSpots in the cirrhotic liver sample (Fibrotic_4) were located in severely fibrotic areas of the tissue that could not be further histologically categorized.

Although overall SenSpot abundance scaled with disease progression,^12^ the proportion of SenSpots to non-senescent tissue was non-linear across stages (**Extended Data Figure 4F**), because gene expression was normalized within each tissue sample. This approach isolated the top M4M-scoring microenvironments and prioritized capturing peak, localized senescence. Top-scoring SenSpots peaked in density during early inflammation (Inflammation_1), reinforcing the concept that focal inflammatory stress serves as a primary trigger for the development of fibrotic niches. Tissue-wide zonation changes with disease progression were further investigated using the percent of total Visium spots containing periportal or pericentral hepatocyte gene signatures. While zonation proportions remained stable across early disease stages, advanced cirrhosis exhibited a marked loss of pericentral cell identity accompanied by an expansion of periportal-type identity (**Extended Data Figure 4G**).

### Spatial Deconvolution and Pathway Analysis

To identify the spatial distribution of gene expression networks in the liver tissue, spatial deconvolution^19^ was performed and identified 14 topics of co-expressed genes across all eight tissue samples. **Figure 3A** provides the spatial map of the 14 topics across each tissue. A SenSpot ratio (SR) quantifying the strength of association for each deconvolution topic with senescent niches identified Topics 7, 10 and 1 as most highly co-localized within SenSpots (**Figure 3B**, Y-axis ranking). We further applied consensus cell phenotyping markers to estimate the prevalence of different cell types contributing to each topic (**Figure 3B**, X-axis labels and Z-score heatmap). This highlighted that topics 7, 10 and 1 were high in multiple immune cells including B cells, dendritic cells (DC), Kupffer cells, monocytes, neutrophils, and T cells. These topics were also low in sinusoidal endothelium, and NK cells. Although proportions of all topics were heterogeneous across the liver samples (**Figure 3C**), the SenSpot-associated topics (represented by colors orange, red, and dark pink, respectively) demonstrated specific enrichment patterns (**Figure 3D**). Topic 1 was proportionally elevated in two of the normal tissues and the steatosis_1 sample. Topic 7 and Topic 10 were found in all tissues, but were a dominant feature of the cirrhotic sample, Fibrotic_4.

**Figure 3.**
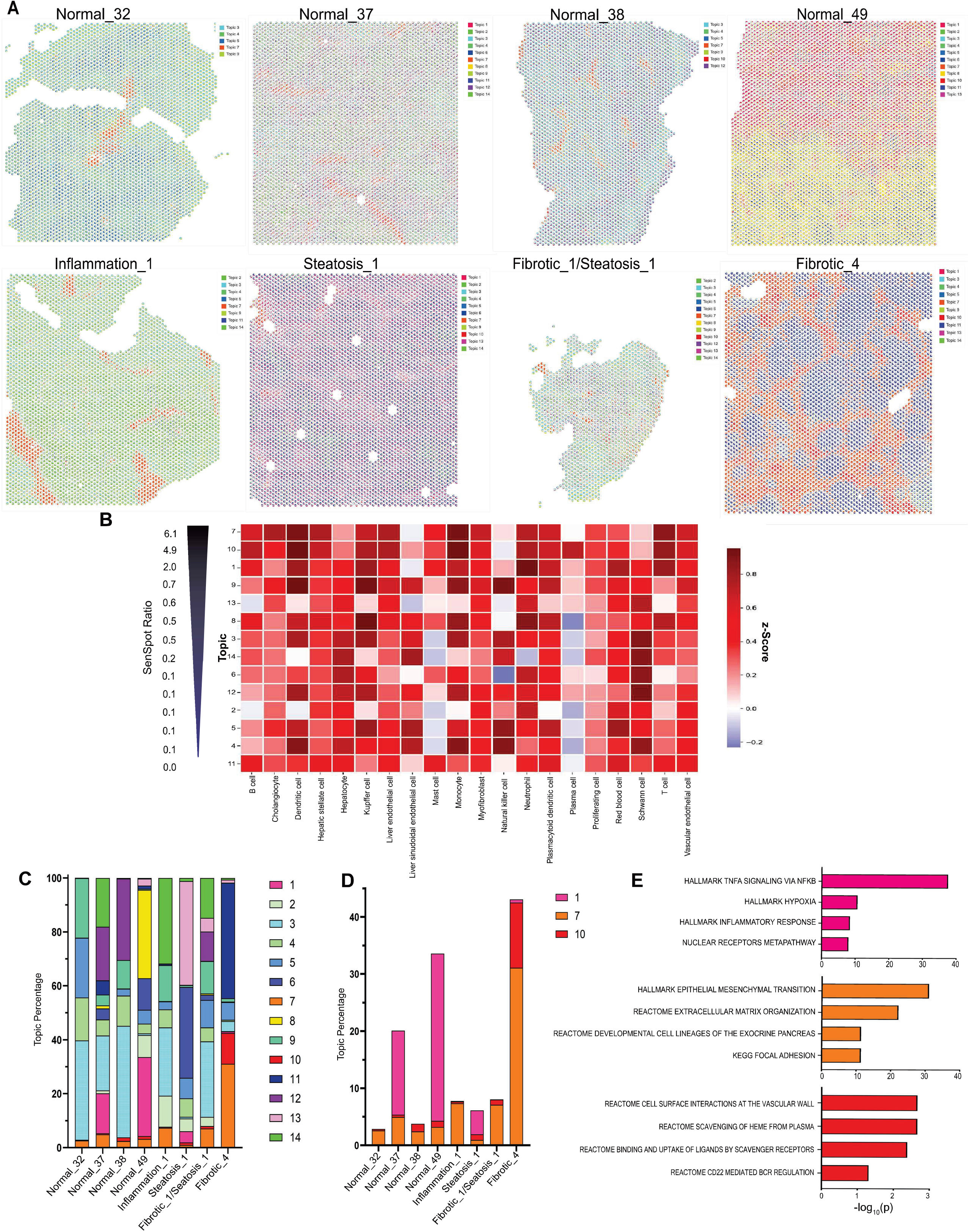
Deconvolution identifies cell types and pathways enriched in SenSpots. **A.** Spatial distribution of the 14 deconvolution topics across all 8 liver samples. **B.** Heat map of topics 1-14 (rows) across liver cell types (columns), ordered top to bottom by abundance of SenSpots in each topic based on the SenSpot Ratio (left arrow). **C.** Percentage of each topic represented in each liver sample. **D.** Percent of SenSpot-enriched topics number 1 (pink), 7 (orange), and 10 (red) in each liver sample. **E.** Pathway analysis of the topmost upregulated genes in SenSpot enriched topics number 1 (pink), 7 (orange), and 10 (red), negative log transformed adjusted p-values.

Pathway enrichment analysis of the SenSpot dense topics 1, 7 and 10 was performed (**Figure 3E**). Topic 1 was predominated by TNFα signaling via NF-κB activation, driven by genes such as *NFKB1/2* and *CCND1*, and was associated with liver cell survival and regeneration.^20^ Topic 7 had epithelial to mesenchymal transition (EMT) and ECM organization pathways driven by genes such as collagens, *ACTA2*, *FBLN1/2*, and *ITGA7*. Three out of four top upregulated pathways in Topic 10 were driven by genes involved in the complement pathway including *COLEC11*, *IGKC*, and *IGLC1*, and are associated with immune activation, apoptosis and cell debris clearance. In summary, spatial deconvolution identified three topics of co-expressed genes significantly enriched within SenSpots (Topics 1, 7, and 10). Topic 1, characterized by genes driving liver regeneration, was strongly represented in healthy SenSpots and early steatosis (Steatosis_1), but was strikingly absent in three of four diseased tissues. Conversely, Topic 7 (EMT and ECM organization) and Topic 10 (complement and B-cell activation) were present across all stages, yet reached peak abundance in advanced cirrhosis, highlighting a central role in disease progression.

### Spatial Signal Expansion (SSE) surrounding SenSpots increases with fibrotic disease

To investigate transcriptional regulation of key pathways in the microenvironment surrounding the SenSpots, we developed an approach termed Spatial Signal Expansion (SSE). **Figure 4A** illustrates the premise of SSE for investigating gene expression in the SenSpot microenvironment from distance 0, the SenSpot, to a distance of 5 Visium spots away, covering approximately 500 µm of surrounding tissue. This method identifies whether a gene expression program is upregulated in SenSpots and rapidly normalizes to baseline or extends outward into the adjacent microenvironment. The SSE approach identified strong localized upregulation of the M4M gene categories (cell stress, SASP, anti-apoptotic, and ECM remodeling) in the SenSpots with rapid normalization of each module’s mean Z-score to baseline in the immediately adjacent microenvironment across all 8 liver samples (**Figure 4B**). The peak in mean z-score at distance 0 (SenSpot) with rapid normalization to baseline confirmed the unique upregulation of the multiple senescence categories in the SenSpots compared to the surrounding area.

**Figure 4.**
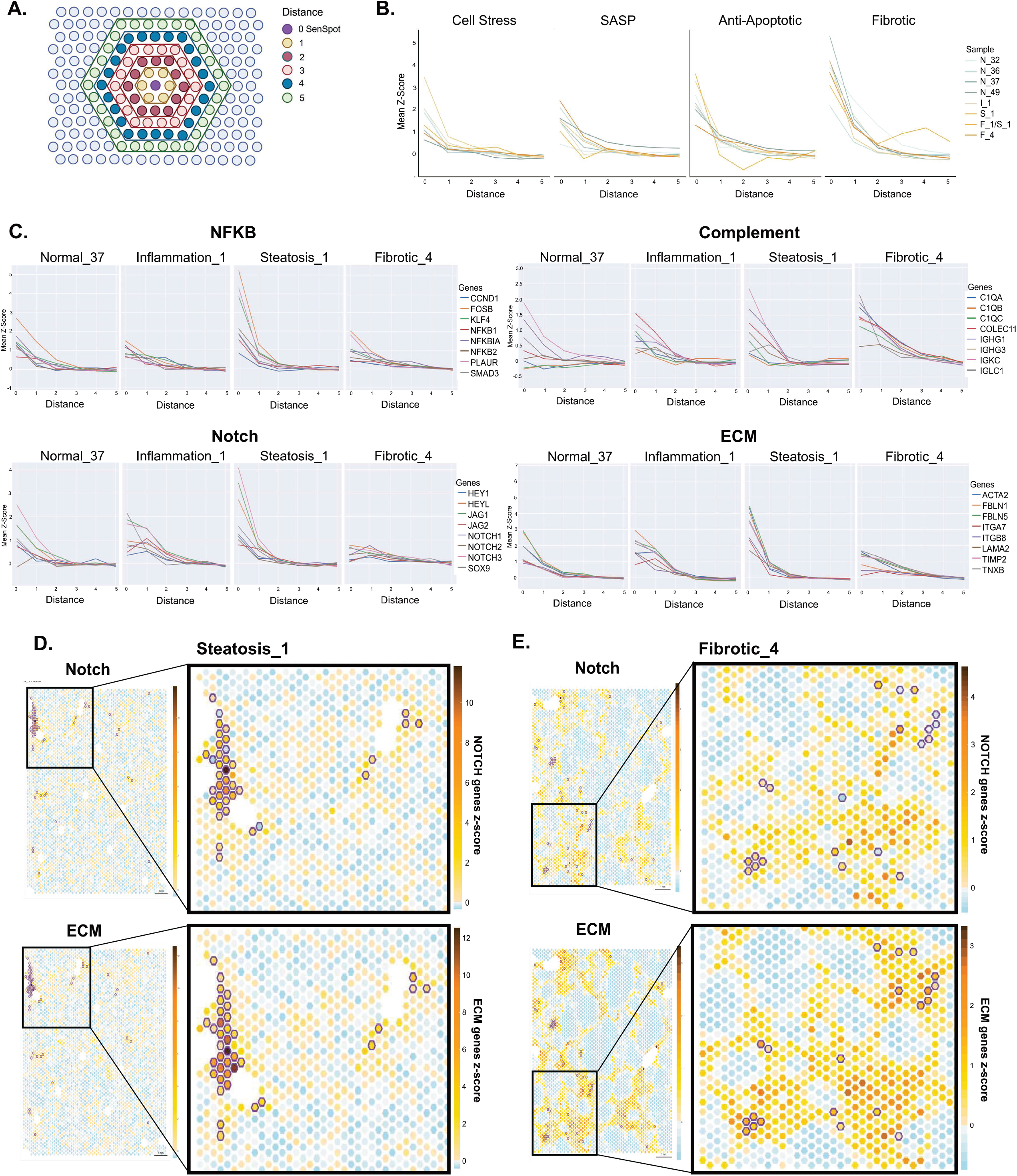
SenSpot microenvironment alterations with disease progression. **A.** Spatial Signal Expansion (SSE) analysis used to investigate the impact of SenSpots on their tissue microenvironment. SSE position 0 is the SenSpot (purple) and each successive Visium spot distance away from the SenSpot are distances 1-5. **B.** SSE graphs of the four senescence M4M categories illustrating the mean z-score at distance 0-5 for each liver sample. **C.** SSE graphs of SenSpot enriched pathways, NF-κB, complement, Notch signaling, and Extracellular Matrix (ECM). **D-E.** Spatial heatmaps for Notch signaling and ECM genes for two diseased liver samples. SenSpots are circled in purple. To the right is an enlargement of the tissue area within the black box.

SSE was next utilized to investigate the key pathways identified through DEG characterization and spatial deconvolution of the SenSpots: NF-κB and NOTCH signaling, complement activation and EMT/ECM remodeling (**Figure 4C**). SSE plots for individual samples demonstrate changes in spatial expansion patterns of pathway genes across representative samples of the disease spectrum in this cohort (**Figure 4C**, **Supplemental Figure 1**).

NF-κB showed peak upregulation within SenSpots and rapid normalization toward baseline between distances 1 and 2 for the healthy tissue samples and was most dramatically exaggerated in the steatosis samples. In contrast, the NF-κB SSE graphs for Inflammation_1 and Fibrotic_4 showed lower relative upregulation within the SenSpots themselves and a more gradual return to baseline in the surrounding distances, suggesting a more generalized level of NF-κB activity outside of the senescent niches in these disease states.

Individual genes within the complement pathway showed divergent SSE patterns in normal samples. Complement C1Q complex genes (*C1QA*, *C1QB* and *C1QC*), responsible for initiating apoptotic cell elimination and clearing of cellular debris were upregulated only in diseased tissues. On the other hand, *IGHG1*, *IGHG3*, *IGKC*, and *IGLC1* (components of antibody structure) were upregulated in all SenSpots indicating the presence of plasma cells.^21^ Fibrotic_4 SenSpots had the greatest upregulation of genes in the complement pathway, with a more gradual return to baseline at distance=3.

The NOTCH and ECM pathway genes were upregulated in SenSpots from normal and steatosis_1 tissue samples with rapid normalization toward baseline between distances 1 and 2. In contrast, the inflammation_1 sample showed more gradual declines in these pathways, consistent with expansion of these processes outside of the localized senescent niche. The Fibrotic_4 sample showed low peak ECM enrichment in SenSpots with very gradual normalization, consistent with the widespread fibrosis. Fibrotic_4 also showed almost no enrichment for reparative NOTCH signaling within the SenSpots, illustrated by a mean Z-score <1 at distance zero and essentially flat SSE curves extending over distance.

The spatial heatmaps of SSE in **Figure 4D** and **Supplemental Figure 2** allow further interpretation of the expression of senescence features. In Fibrotic_4, the SenSpots showed low to moderate expression of NOTCH signaling gens while closely adjacent spots frequently had moderate to high expression. Similarly, ECM pathway z-scores were moderate within SenSpots, but frequently high in immediately adjacent areas. Therefore, the flat SSE curves for NOTCH, ECM and NF-κB in Fibrotic_4 represent a dispersed expression of these pathways expanding into surrounding areas.

In summary, SSE profiling of the SenSpot microenvironment validated the localized enrichment of core SenSpot associated pathways uncovering distinct spatial behavior across disease stages. NF-κB, complement, NOTCH, and ECM pathway signals exhibited localized SenSpot enrichment in normal and mild pathology tissues, whereas advanced cirrhosis revealed a broad spread of pro-inflammatory and fibrotic signaling throughout the surrounding tissue microenvironment.

### scRNAseq corroborates senescent cell lineages and pathway crosstalk identified in SenSpots

To validate the cell-types and functional programs operating within SenSpot microenvironments, we performed scRNA-seq on 24 human liver biopsies spanning diverse stages of liver pathology (**Figure 5A**). Tissues were processed using a probe-based scRNAseq workflow following fixation and dissociation, yielding consistent cell-type clustering and representation across all samples (**Extended Data Figures 5A–B**).

**Figure 5.**
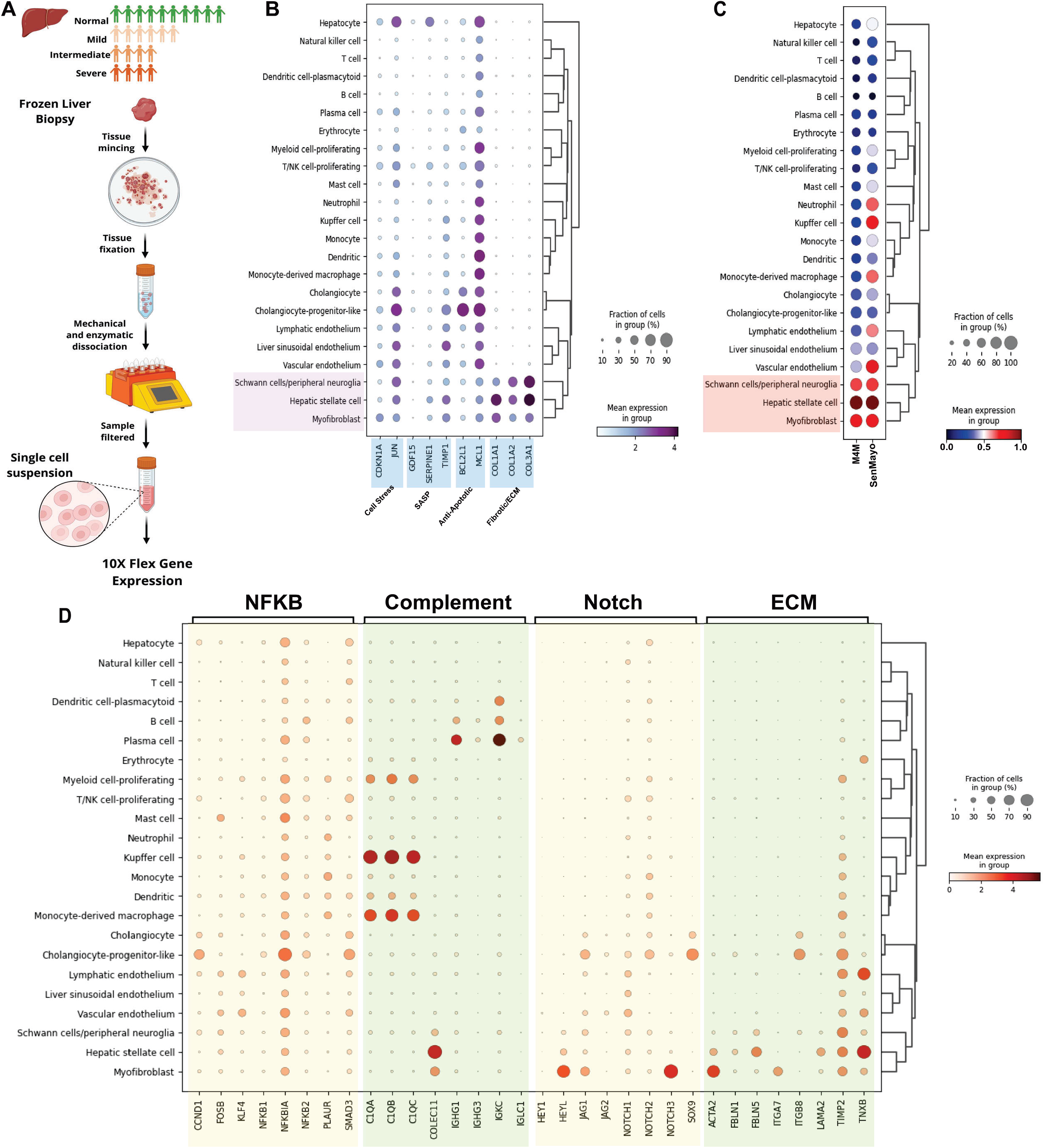
Single-Cell RNA sequencing corroborates senescent cell lineages and pathway crosstalk identified from SenSpots. **A.** 24 frozen liver biopsies graded as normal, mild, intermediate or severe pathology were processed for 10x Genomics Flex Gene Expression profiling. **B**. Expression of the M4M genes by cell-type for all of the samples combined. **C.** Expression of M4M and SenMayo gene sets summarized as an average module score per cell-type. Values are scaled between 0 and 1 to highlight the cell-types with highest senescence scores according to both gene sets in the combined samples. **D.** Expression of key genes identified in the spatial analysis in the scRNAseq datasets.

Mapping our 10-gene M4M gene panel across the single-cell transcriptomes identified Schwann cells, hepatic stellate cells (HSCs), and myofibroblasts as the primary drivers of hepatic senescence, exhibiting concurrent expression across all four M4M functional categories (**Figure 5B**). In contrast, other lineages (*i.e.,* hepatocytes, cholangiocytes and progenitor-like cells) were enriched in three categories, but lacked the ECM gene module. To summarize the expression of senescence markers with a single metric, we calculated a module score comparing each cell gene expression pattern to the tissue average using our M4M panel and benchmarked it against the SenMayo signature.^22^ Schwann cells, HSCs, and myofibroblasts maintained high mean senescence feature expression across both gene sets (**Figure 5C**). The SenMayo gene set, developed in the absence of cell cycle inhibitor expression, additionally identified neutrophils, Kupffer cells, monocyte-derived macrophages, and endothelial sub-populations, driven primarily by the broad inclusion of SASP components in the gene set.

To identify which cell-types drive the pathways identified by spatial deconvolution, we queried the single-cell dataset with our pathway-specific gene sets (**Figure 5D**). NF-κB signaling was enriched in mast cells, cholangiocyte progenitor-like cells, and vascular/lymphatic endothelial cells. Complement pathway C1q complex genes (*C1QA*, *C1QB*, *C1QC)* were predominantly expressed by monocyte-derived macrophages, Kupffer cells, proliferating myeloid cells, and dendritic cells. *COLEC11* expression was restricted to the core senescent stromal triad (Schwann cells, HSCs, myofibroblasts), whereas immunoglobulin genes (*IGHG1*, *IGHG3*, *IGKC)* were localized to plasma cells, B cells, and plasmacytoid dendritic cells. NOTCH signaling was heavily driven by myofibroblasts and cholangiocyte progenitor-like cells. ECM-remodeling genes were co-expressed by myofibroblasts, HSCs, Schwann cells, endothelial lineages, and progenitor-like cholangiocytes.

To explore whether these top senescent cell lineages scale with clinical pathology, we quantified their expression across disease stages (**Extended Data Figure 5C-F**). This confirmed a progressive, disease-stage-dependent expansion of senescent cell populations in HSCs, myofibroblasts, and Schwann cells with increases in cell stress and SASP gene expression with disease progression (**Extended Data Figure 5C**). Multiple liver cell types, including epithelial and immune cell lineages revealed increases in M4M gene expression with disease progression in the cell stress, SASP and anti-apoptotic categories (**Extended Data Figure 5D-F**), most notably in cholangiocyte-progenitor-like cells, hepatocytes, and proliferating T and NK cells.

In summary, scRNAseq identified Schwann cells, hepatic stellate cells, and myofibroblasts as the lineages with the highest expression of M4M-integreated senescence markers in human liver disease, while mapping distinct paracrine divisions of labor ranging from C1q-mediated macrophage activity to endothelial and progenitor NF-κB activation that progressively expanded from SenSpots with advancing pathology. The most marked changes in gene expression profiles occurred in immune subpopulations and cholangiocyte-progenitor-like cells.

### NicheNet modeling resolves multi-cellular paracrine networks and senescence signals driving the fibrotic niche

Given the complex multicellular changes that accompanied disease progression to cirrhosis, we next sought to identify intercellular signaling networks that drive the senescence-associated phenotypes observed in diseased tissue. The NicheNet pipeline was applied to our scRNA-seq datasets to infer ligand–receptor interactions and their downstream transcriptional consequences in receiver cells. Unlike approaches based solely on ligand– receptor co-expression, NicheNet prioritizes signaling interactions based on their ability to predict transcriptional changes within receiver cells,^23^ allowing us to infer and compare paracrine signaling networks between normal and fibrotic liver tissues. We interrogated (i) which ligands from sender cells in fibrotic livers predict transcriptional alterations in receiver cells, (ii) which differentially upregulated receptors in receiver cells can interact with these predicted ligands, and (iii) which DEGs are predicted downstream targets of these receptors (**Figure 6A-C**).

**Figure 6:**
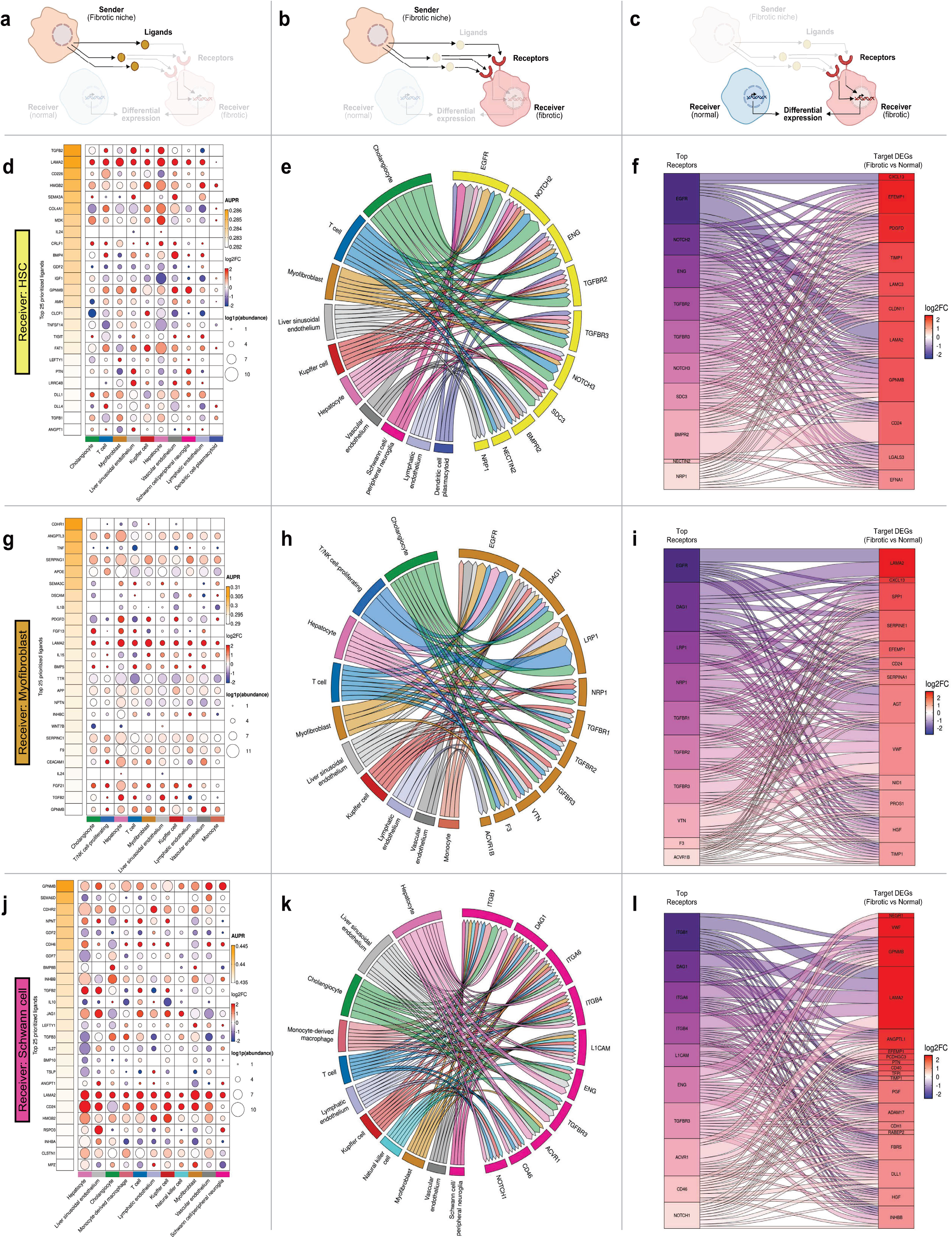
NicheNet modeling identifies multicellular paracrine signaling networks associated with senescent stromal populations in fibrotic liver. A-C. Schematic of the NicheNet analysis workflow used to interrogate cell-cell communication in fibrotic vs normal liver samples. A. Ligands expressed by potential sender cell populations were prioritized based on their ability to predict differential gene expression in the indicated receiver population. B. Differentially upregulated receptors in receiver cells were mapped to ligands identified in A. C. Differentially expressed genes in receiver cells were subsequently linked to upstream receptors to identify predicted downstream transcriptional targets. D-F. NicheNet analysis of hepatic stellate cells (HSCs) as receiver cells. D. Dot plot showing predicted ligands upregulated across sender cell-types. Dot size represents ligand expression level and the color indicates log2 fold-change within the sender cell-type in fibrotic vs normal livers. The ligands were ranked by area under the precision-recall curve (AUPR). E. Circos plot showing the predicted interactions between sender cell-types and differentially upregulated receptors in receiver cells. The width of connecting ribbons correlate to weighted contribution of each sender cell to individual ligand-receptor interactions. F. Sankey plot linking prioritized HSC receptors to predicted downstream differentially expressed target genes within receiver cells, colored by log2 fold-change in fibrotic vs normal livers. G-I. Ligand prioritization, sender-receptor interactions, and receptor-target relationships for myofibroblasts and J-L. Schwann cells as receiver cells.

We selected the three stromal cell types: hepatic stellate cells (HSCs), myofibroblasts, and Schwann cells, which exhibited the highest expression of senescence-associated signatures in our single-cell analysis, as receiver populations to assess how microenvironment inputs might impact these cell types central to the observed senotype (**Figure 6D-L**, **Table 1**). Across these three populations, predicted signaling was highly interconnected, with similar ligand signatures and sender cell-types detected. Many of the top shared ligands represented ECM components (*LAMA2* and collagens); TGF-β/BMP family proteins; inflammatory and SASP-associated factors (*TNF*- and *IL*-family members); and NOTCH ligands (*JAG*/*DLL*) (**Figure 6D, G, J**). We next examined the predicted cellular sources of these signals, and cholangiocytes together with hepatocytes, endothelial cells, and immune subsets such as T and NK cells, represented the prominent sender populations in fibrotic livers based on ligand expression and differential regulation (**Figure 6E, H, K**).

**Table 1.** Comparison of Stromal Triad Cell Types as Receivers in Fibrotic Liver.

|  | **Hepatic Stellate Cells (HSCs)** | **Myofibroblasts** | **Schwann Cells / Peripheral Neuroglia** |
|---|---|---|---|
| **Primary Functional Role** | Early transdifferentiation responder, inflammatory cell recruitment, pericyte signaling | Hyper-synthetic ECM deposition, scar contraction, vascular remodeling | Neural guidance, axon sprouting, basement membrane anchorage, neuro-vascular support |
| **Sender Cells** | Cholangiocytes, T-Cells, Myofibroblast, LSECs, Kupffer Cells, Schwann Cells | Cholangiocytes, T/NK Cell-Proliferating, Hepatocytes, T-Cells, Myofibroblasts, LSECs | Hepatocytes, LSECs, Cholangiocytes, Macrophages/Kupffer Cells, Myofibroblasts, Schwann Cells |
| **Ligands Received** | TGFB2, LAMA2, HMGB2, SEMA3A, COLA41, MDK, GPNMB, FAT1, DLL1/4, ANGPT1 | ANGPTL3, TNF, SERPING1, APOE, SEMA3C, PDGFD, FGF13/21, LAMA2, BMP5, TGFB2, GPNMB | GPNMB, SEMA6D, CDHR2, NPNT, INHBB, TGFB2, JAG1, LAMA2, CD24, HMGB2 |
| **Receptors** | EGFR, NOTCH2/3, ENG, TGFBR2/3, SDC3, BMPR2, NECTIN2, NRP1 | EGFR, DAG1, LRP1, NRP1, TGFBR1/2/3, VTN, F3, ACVR1B | ITGB1/4, DAG1, ITGA6, L1CAM, ENG, TGFBR3, ACVR1, CD46, NOTCH1 |
| **Upregulated Genes in recipient cells** | CXCL13, EFEMP1, PDGFD, TIMP1, LAMC3, CLDN11, LAMA2, GPNMB, CD24, LGALS3, EFNA1 | LAMA2, CXCL13, SPP1, SERPINE1, EFEMP1, CD24, SERPINA1, AGT, VWF, NID1, PROS1, HGF, TIMP1 | NEGR1, VWF, GPNMB, LAMA2, ANGPTL1, EFEMP1, PCDHGC3, PTN, CD40, TFP1, TIMP1, PGF, ADAM17, CDH1, RABEP2, FBRS, DLL1, HGF, INHBB |

Red represents genes identified in at least 2/3 stromal triad cells

Extending this analysis across the broader single-cell dataset revealed that these communication networks were highly interconnected (**Extended Data Figure 6**). For example, although T and NK cells contributed inflammatory signals to the fibrotic niche, predicted signaling to these cells suggested that inflammatory cues may originate from cholangiocytes, myofibroblasts, hepatocytes, endothelial and immunes subsets (**Extended Data Figure 7-8)**. Thus, the senescence-associated fibrotic niche appears to emerge from coordinated paracrine communication across multiple cell populations, supporting a multicellular model of stromal niche regulation in fibrotic liver.

The stromal triad cells (HSCs, myofibroblasts, and Schwann cells) exhibited overlapping receptor and downstream transcriptional responses in fibrotic disease. All three cell types shared upregulation of TGF-β/Activin pathway receptors (*TGFBR1*/*2*/*3*, *ACVR1B*, *ACVR1*) and NOTCH pathway receptors (*NOTCH1*/*2*/*3*, *SDC3*, *LRP1*, and *NRP1*). HSCs and myofibroblasts shared *EGFR* receptor expression and myofibroblasts and Schwann cells shared upregulation of the adhesion receptor *DAG1*. Schwann cells additionally expressed a distinct receptor profile enriched with integrins receptors that are drivers of hepatocellular carcinoma (*ITGB1*/*4*, *ITGA6*)^24,25^ (**Figure 6E,H,K**, **Table 1**).

These receptor configurations reflect stress-associated activation and matrix-remodeling programs within fibrotic tissue, a finding substantiated by predicted downstream target gene activation across all three receiver cell types. Specifically, NicheNet signaling converged on a universal downstream matrix-remodeling core comprising *LAMA2*, *TIMP1* and *EFEMP1* (**Figure 6F,I,L, and Table1**). Lineage-specific downstream outputs further revealed that HSCs and myofibroblasts activated *CXCL13* (a potent B-cell chemoattractant)^26,27^ and *CD24* (an anti-phagocytic “don’t eat me” signal),^28,29^ whereas *GPNMB*, *VWF* and *HGF* were upregulated in two receiver cells (Schwann cells in combination with either HSC or myofibroblasts) (**Figure 6F,I,L**). The common upregulated genes (*GPNMB*, *VWF* and *HGF*) were all associated with liver disease progression and identified as potential therapeutic targets.^27,30–34^ Collectively, these findings demonstrate that senescent stromal cells function as integral nodes within a highly interconnected multicellular signaling network that actively structures the fibrotic liver niche (**Extended Data Figure 9**).

## Discussion

Cellular senescence is a complex, heterogeneous biological state characterized by stable cell-cycle arrest, resistance to apoptosis, metabolic alterations, and the secretion of a pro-inflammatory senescent-associated secretory phenotype (SASP). Because no single biomarker uniquely defines senescent cells *in vivo*, spatial transcriptomic identification of senescence niches in human disease has remained a major challenge. In this study, we addressed this complexity by developing “SenSpots,” a novel, multi-attribute spatial framework integrating cell stress (*CDKN1A*, *JUN*), SASP signaling (*GDF15*, *SERPINE1*, *TIMP1*), anti-apoptotic survival pathways (*BCL2L1*, *MCL1*), and ECM remodeling (*COL1A1*, *COL1A2*, *COL3A1*). By combining spatial transcriptomics across human liver biopsies spanning healthy to cirrhotic with scRNA-seq, we established high-resolution mapping of hepatic senescence. Methodologically, integrating spatial transcriptomics with scRNAseq bridges spatial coverage and cellular resolution, while rigorous artifact filtering overcomes tissue-edge effects particularly inherent to smaller liver biopsies. This approach uncovered a fundamental functional distinction between the beneficial, transient senescence during physiological tissue repair observed in non-pathologic samples and the toxic, chronic senescence observed in liver samples with fibrosis progression. Interestingly, these results identify senescent Schwann cells as a novel cellular driver within the hepatic niche. Our findings revealed distinct biologic programs in senescent niches in samples with MASLD progression. Although SenSpots across all stages share a core signature of ECM remodeling and calcium-dependent signaling, their biological outputs diverge sharply. In healthy tissue, SenSpots exhibit a restorative, wound-healing phenotype that is consistent with beneficial, transient senescence that restricts acute injury and facilitates repair.^35^ In contrast, fibrotic SenSpots are persistent, maladaptive niches dominated by chronic inflammation, active ECM disassembly, and TGFβ-1-driven paracrine transmission.^11,36^

The spatial localization of SenSpots to portal tracts redefines our understanding of how cellular senescence intersects with the liver’s intrinsic regenerative biology. Portal tracts are recognized as critical anatomical niches housing the liver progenitor cell pool and orchestrating repair following chronic injury.^37,38^ Our findings indicate that early in MASLD, cellular senescence is an active component of the portal progenitor/ cholangiocyte response (marked by *SOX9* and *EPCAM* activation).^39–41^ While transient activation of NOTCH and C1q signaling within these early niches mirrors canonical programs of controlled tissue repair,^39^ progressive fibrogenesis disrupts this localized equilibrium. As disease advances toward cirrhosis, the well-established paradigm of metabolic zonation collapses, coinciding with the breakdown of pro-regenerative signaling cascades and the spatial consolidation of senescent cells into expanding fibrotic septa. Furthermore, our Spatial Signal Expansion (SSE) analysis provides a mechanism for mapping the extent of the “paracrine field effect” in chronic liver disease.^42^ While early senescent signaling remains strictly compartmentalized within the portal triad, advanced cirrhosis transitions to homogeneous parenchymal dissemination of NF-*κ*B, NOTCH, and ECM remodeling pathways, effectively transforming a localized repair response into a tissue-wide driver of disease progression.

By resolving the multicellular composition within the senescence niches, this study extends the classical view of the fibrotic liver stroma. While hepatic stellate cells (HSCs) and myofibroblasts have been established as primary drivers of extracellular matrix deposition,^14,43^ our integration of spatial and single-cell transcriptomics uncovers senescent Schwann cells as an unaccounted-for stromal lineage in human fibrotic disease. Our data support a model where a senescent stromal triad comprising HSCs, myofibroblasts, and Schwann cells work cooperatively within the senescent niche to expand the fibrotic septa (**Extended Data Figure 9**). By identifying cholangiocytes as primary signaling senders targeting the stromal triad, these data provide a spatial and cellular basis for how biliary stress coordinates surrounding stromal remodeling. Rather than functioning as a non-specific scar, this reactive stroma executes a specialized matrix program that simultaneously accelerates basement membrane assembly and inhibits metalloproteinase-mediated turnover.

Furthermore, the convergence of activated HSCs and myofibroblasts on hyperactive EGFR signaling bridges chronic fibrogenesis with pre-malignant transformation. EGFR hyperactivation is a recognized catalyst of MASH progression^44^ and a potent driver of senescence-associated therapeutic resistance and oncogenesis.^45–48^ By placing EGFR signaling within the senescent stromal niche, our findings link persistent fibrotic remodeling to the microenvironmental priming required for hepatocellular carcinoma (HCC) development. This pathogenic stroma is further reinforced by localized immune-evasive cross-talk: stromal upregulation of CXCL13 provides a mechanism for pathogenic B-cell recruitment that dampens regenerative capacity,^26,27^ while CD24 expression presents a clearance barrier that impairs macrophage-mediated phagocytosis of senescent cells,^28,29^ effectively perpetuating the chronic fibrotic niche.

Located within the portal nerve plexus, senescent Schwann cells express high baseline levels across all four M4M functional modules, establishing these portal neuroglial cells as previously unappreciated drivers of neuro-vascular and stroma-immune crosstalk during fibrotic remodeling. Schwann cells play an essential role in the protection and regeneration of hepatic nerves. However, sympathetic innervation becomes disorganized and progressively degenerates as MASH/MASLD advances^49^ and the peripheral nervous system damage can further activate the complement pathway.^50^ This neural impairment carries significant functional consequences: chronic stress can disrupt central brain-to-liver signaling and impair liver regeneration via hepatic sympathetic nerve dysfunction,^51^ while neural-lymphatic dysfunction impairs debris clearance and accelerates liver fibrosis.^52^ Together, these findings highlight the critical role of hepatic innervation and demonstrates how progressive neural degeneration accelerates disease progression in the fibrotic liver microenvironment.

In conclusion, this study maps the spatial transition from transient, portal-bounded repair niches to chronic, paracrine-spreading fibrotic fields, uncovering senescent Schwann cells as a novel therapeutic target for senolytic interventions in chronic liver disease.

## Methods

### Human liver tissue procurement, processing, and grading

Human liver tissue was collected in accordance with protocols approved by the University of Minnesota Institution Review Board (STUDY00013764 and STUDY00021098) and through the University of Minnesota human tissue biorepository (BioNet). Histological processing including formalin fixation, paraffin embedding, OCT embedding, sectioning, and histological staining were performed by Biospecimen and Laboratory Services at the University of Minnesota. Hematoxylin and eosin (H&E) and Masson’s trichrome stained tissue slides were used for pathology grading as previously described.^53,54^

### Visium CytAssist spatial transcriptomics

Visium CytAssist for FFPE Spatial Gene Expression 6.5 mm, Human, 4 reaction kit (10x Genomics 1000520), with Visium Human Transcriptome Probe Set v2.0 targeting 18,085 genes with 53,504 probes, was used for liver spatial transcriptomics. A brief overview of Visium CytAssist is in Figure 1A. The following Visium CytAssist spatial transcriptomics protocols were used: 10x Genomics Visium CytAssist Spatial Gene Expression for FFPE - Tissue Preparation Guide (CG000518, Rev B), 10x Genomics Visium v2 CytAssist FFPE Deparaffinization, H&E staining, and imaging (CG000520, Rev B) and Visium v2 CytAssist FFPE Library Construction (CG000495, Rev B/C). Formalin-fixed, paraffin-embedded (FFPE) liver sections of 5 µm thickness were used for each sample. Prior to performing CytAssist-enabled probe transfer, the H&E-stained slides were scanned at 20X with Brightfield on a Huron TissueScope LE at the University Imaging Centers at the University of Minnesota (**Figure 1D, Supplemental Figures 2-3**). Sequencing and 10x Genomics Space Ranger alignment was performed by the University of Minnesota Genomics Center. The Space Ranger pipeline quality and analysis outputs can be found in **Supplemental Table 1** and **Supplemental Figure 1**.

### Artifact removal

BLADE software was used to detect gene expression artifacts attributed to tissue or capture area boundaries and remove the artifactual spots from analysis, as previously described.^55^

### SenSpot identification

The 10 SenSpot genes identified were *CDKN1A* and *JUN* (cell stress); *GDF15*, *SERPINE1*, and *TIMP1* (SASP); *BCL2L1*, *MCL1* (anti-apoptotic); and *COL1A1*, *COL1A2* and *COL3A1* (fibrotic). To ensure contribution of genes from each category in identifying SenSpots, we utilized the mean z-score of all 4 categories, or the mean of four means (M4M) with a threshold of >1.65 (**Figure 1C**).

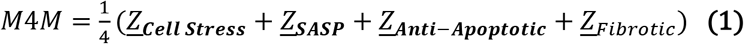

Where:

- 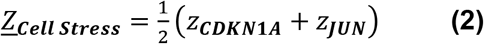
- 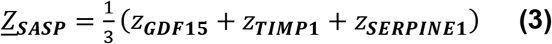
- 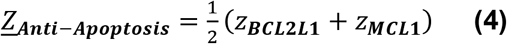
- 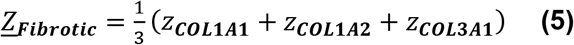

### Differentially expressed genes bioinformatic analysis

Differentially expressed genes (DEGs) in SenSpots Z-score means from the four healthy liver samples (Normal_32, Normal_37, Normal_38 and Normal_49) were used to calculate Ave_Normal.

For interaction network construction, sets of gene identifiers as DEGs were used as node inputs in the STRING interaction interface v12.0 (https://string-db.org). Results were restricted to *H. sapiens* and query gene list only. Molecular interaction modes (edges) with highest confidence score (>0.90) were filtered. Subsequently, nodes and edges were imported to CytoScape (v3.10.0) using the stringApp (v1.7.0) add-on. Nodes were visualized using yFiles layout algorithm and clustered using clusterMaker2 (v2.0) MCL: Markov Cluster Algorithm unsupervised clustering with a granularity parameter of 2.5 for all networks.

Functional enrichment analysis was performed on the gene lists containing the top DEGs. The StringDB 12.0 interface was used, specifying *H. sapiens* as the source organism, filtering criteria: False Discovery Rate (FDR) <0.05, network enrichment signal [weighted harmonic mean between the observed/expected ratio and - log(FDR)] >0.1, and enrichment strength >0.01. The resulting terms were filtered for GO Biological Process (GO:BP) and GO Molecular Function (GO:MF) categories, and a minimum observed gene count of 4 was required for enriched terms. The terms were sorted by signal score. Bubble size was defined by binning the observed gene counts. The plot was generated using pandas, numpy, and matplotlib packages in Python 3.12.11.

For interaction network construction, sets of gene identifiers for DEGs were used as node inputs in the STRING interaction interface v12.0 (https://string-db.org). Results were restricted to *H. sapiens* and query gene list only. Molecular interaction nodes (edges) with highest confidence score (>0.90) were filtered. Subsequently, nodes and edges were imported to CytoScape (v3.10.0) using the stringApp (v1.7.0) add-on. Nodes were visualized using yFiles layout algorithm and clustered using clusterMaker2 (v2.0) MCL clustering with a granularity parameter of 4.0 for all networks.

A heatmap of the mean empirical z-scores across the liver fibrotic disease spectrum was created from DEGs up in Fibrotic_4 after filtering for Ave_Normal. Genes/rows were hierarchically clustered by Euclidean distance and average linkage then plotted using the pandas, numpy, matplotlib, seaborn and scipy packages in Python 3.12.11.

Spatial heatmaps and the deconvolution topic heatmaps are products of custom software (see the libraries HeatMap.py and ScatterPie.py in the code repository at https://github.com/KummerfeldLab/2025-Visium-CytAssist-liver-paper) written in Python utilizing Bokeh https://docs.bokeh.org figure rendering functions and Seaborn https://seaborn.pydata.org color manipulation routines.

### Pathology SenSpot annotation

To annotate zonation of SenSpots, loupe Space Ranger output files containing gene expression data were mapped onto histological images opened in Loupe Browser software (version 8.0.0, 10x Genomics). Identified SenSpot barcodes were imported from CSV files, manually annotated by a pathologist based on tissue features, then exported. Spots were given one of 6 labels: Zone 1 - Tract (>50% of the spots were within a portal tract); Zone 1 - Periportal (>50% of the spots were in liver parenchyma within 100 μm of a portal tract boundary); Capsule (>50% of the spots were within the liver capsule layer); Zone 2 (>50% of the spots were within the midlobular parenchyma); Zone 3 (>50% of the spots were within 100 μm of the central vein); or Fibrosis (a spot containing dense fibrosis without definitive zonation).

### Spatial deconvolution

We applied the STDeconvolve algorithm^19^, a reference-free approach to determination of the cell composition within a Visium spot to the aggregated BLADE-filtered sample gene count tables. We used the procedure described here https://jef.works/STdeconvolve/additional_features.html to select the number of topics to learn, arriving at 14 topics to best strike the balance between introducing higher model perplexity, insufficient granularity, and overfitting. Higher topic number models had a higher number of ‘sub-topics,’ small variations in the existing topics that had too small an abundance to be useful. Each topic is a set of estimated ratios of gene counts, which were then used to estimate topic presence ratios for all spots. The presence of each topic at each spot was then visualized.

Pathway analysis was performed using the set of 1,000 most differentially expressed genes across the 14 identified topics, we converted the expression values to per-gene z-scores in R (v.4.4.2). Then genes were filtered to retain only those with an absolute value of the Z-score was >2. Genes meeting this threshold for each topic were used for enrichment analysis by MSigDB Hallmark, KEGG legacy, KEGG medicus, Reactome and Wikipathways. The full set of 1,000 genes in the topic analysis was used as the background for over-representation analysis.

The **SenSpot Ratio** identifies the increase or decrease in the presence of a topic in SenSpots. It was calculated by dividing the mean of the StDeconvolve-modeled percentage of a topic in SenSpots by the same mean from all spots.

### Spatial Signal Expansion

To examine the expansion of pathway expression and senescence affiliated gene groups, the z-score values of each gene in a respective pathway or gene group were averaged at 1 to 5 Visium spots encircling a SenSpot.

### Single-Cell RNA Sequencing (scRNAseq)

24 donors were selected for probe-based single-cell RNA-sequencing analysis, 5 of which are also represented in the Visium datasets (V42L29-278_D1, V42L06-053_D1, V42L29-055_A1, V42L29-055_D1, V42L06-053_A1). Liver samples included those collected prospectively between 2022 and 2023, as well as archived cases collected between 1999 and 2019 and procured from the biorepository at the University of Minnesota’s Clinical and Translational Research Institute. Chromium Fixed RNA Profiling (10x Genomics) enables single-cell transcriptomic analysis of even suboptimal or degraded sources by using short oligonucleotide probe chemistry similar to Visium, representing the same gene set. For 16 retrospective cases, cell suspensions were generated from snap frozen tissues using the tissue fixation and dissociation method specified by the manufacturer (10x Genomics CG00053 Rev A/B). The remaining 8 suspensions were generated from OCT-embedded frozen tissue by modifying the manufacturer’s protocol.

Fixed cell suspensions were hybridized with probes, pooled into groups of 4, captured and barcoded using the ChromiumX instrument, and lysed for sequencing library preparation in 3 different batches (batch 1, n=4; batch 2, n=12, batch 3, n=8) following the manufacturer’s instructions (10x Genomics CG000527 Rev C-E). Libraries were sequenced with paired-end dual-index read format (28/10/10/90) on either an Illumina NovaSeq 6000 S1/S4 flow cell (batch 1 and 2) or an Element AVITI Cloudbreak Freestyle High Output kit (batch 3). Raw base call files were converted to FASTQ format using bcl2fastq software (Illumina) then aligned using the cellranger multi-invocation of 10x Genomics Cell Ranger software (v7.1.0 for batch 1 and 2; v8.0.1 for batch 3). Cells that could not be clearly identified were filtered out of the dataset for downstream analysis, leaving a total of 198,674 cells. The pathological stage of each sample was defined based on steatosis, ballooning, inflammation, and fibrosis scores. “Normal” cases had a total score of zero; “mild” cases had a score of one; “intermediate” cases had a score >1 and a fibrosis score <2; “severe” cases had a score >1 and a fibrosis score >2 (**Supplemental Table 3**).

Senescence scores were calculated using the harmonized gene expression across all samples. To identify the cells with higher expression of senescence gene sets (M4M, SenMayo) in the whole tissue, a general module score was calculated with the tl.score_genes function of the scanpy package. Then, to assess the relationship between senescence and disease progression, scores were calculated for each cell type. The expression levels of each M4M gene were z-scaled using the mean and standard deviation in each cell type. The M4M score was calculated as indicated in section “SenSpot identification” equations (1) - (5) (**SenSpot identification** section). The average score per cell type and sample was calculated using a minimum threshold of 5 cells, and used to fit a linear model (average_senescence_score ∼ pathology stage) where the slope represents the change in M4M expression level with disease progression. Slope estimates significantly different from zero were selected based on the linear model p.values after FDR correction.

### NicheNet ligand-receptor interaction analysis

Intercellular communication was inferred using the NicheNet framework.^23^ The annotated single-cell expression matrix, cell metadata, and UMAP coordinates were imported into R as a Seurat object. Donor age and disease annotations were mapped to individual cells. For each comparison, receiver cell populations and conditions, defined by age and disease groups, were selected. All annotated cell populations were considered potential sending populations. Receiver-cell differentially expressed genes were identified using Seurat’s FindMarkers function, filtering for the fraction of cells expressing the receptor at ≥10% and |log2_FC_| >0.75. NicheNet’s ligand-receptor network and ligand-target matrix were used to identify candidate ligands and predict their regulatory potential. Ligands were ranked by area under the precision-recall curve (AUPR). Candidate ligand-receptor interactions were further evaluated based on ligand and receptor expression in the corresponding sender and receiver populations. For chord diagram visualization, sender ligands were retained when |log2_FC_| >0.75 and log(1+expression) ≥5. Chord diagrams display the top 30 ligands ranked by AUPR and the top 10 prioritized receptors. Receptor-to-target Sankey diagrams linked receptors to candidate ligands and their predicted downstream targets using NicheNet ligand-target regulatory potential. Downstream target genes were filtered using |log2_FC_| >0.75. Up to 10 targets per receptor were ranked according to predicted regulatory potential.

## Acknowledgements

This work was supported in part by the resources and staff at the University of Minnesota Genomics Center [SCR_012413], the University Imaging Centers [SCR_020997] and the University CTSI Histology & Digital Imaging Laboratory [UL1TR002494].

---

**Extended Data Figure 1.**
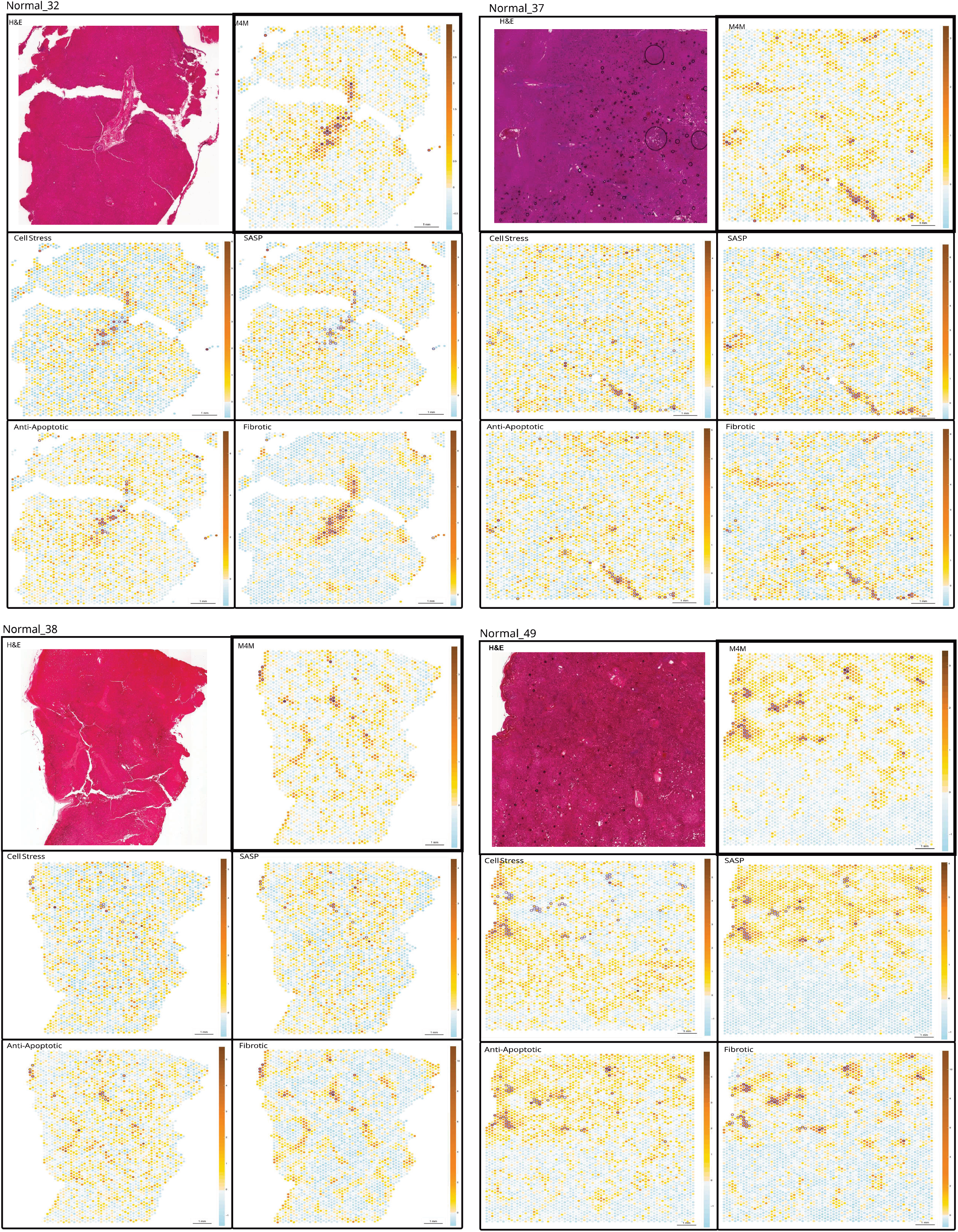
Images of Visium data identifying SenSpots in healthy liver samples. H&E (hematoxylin and eosin) stain of each tissue slide used for Visium, spatial heatmaps for M4M and the mean z-score for each senescence category: cell stress, SASP, anti-apoptotic, and fibrotic. SenSpots identified by M4M are highlighted with purple circles.

**Extended Data Figure 2.**
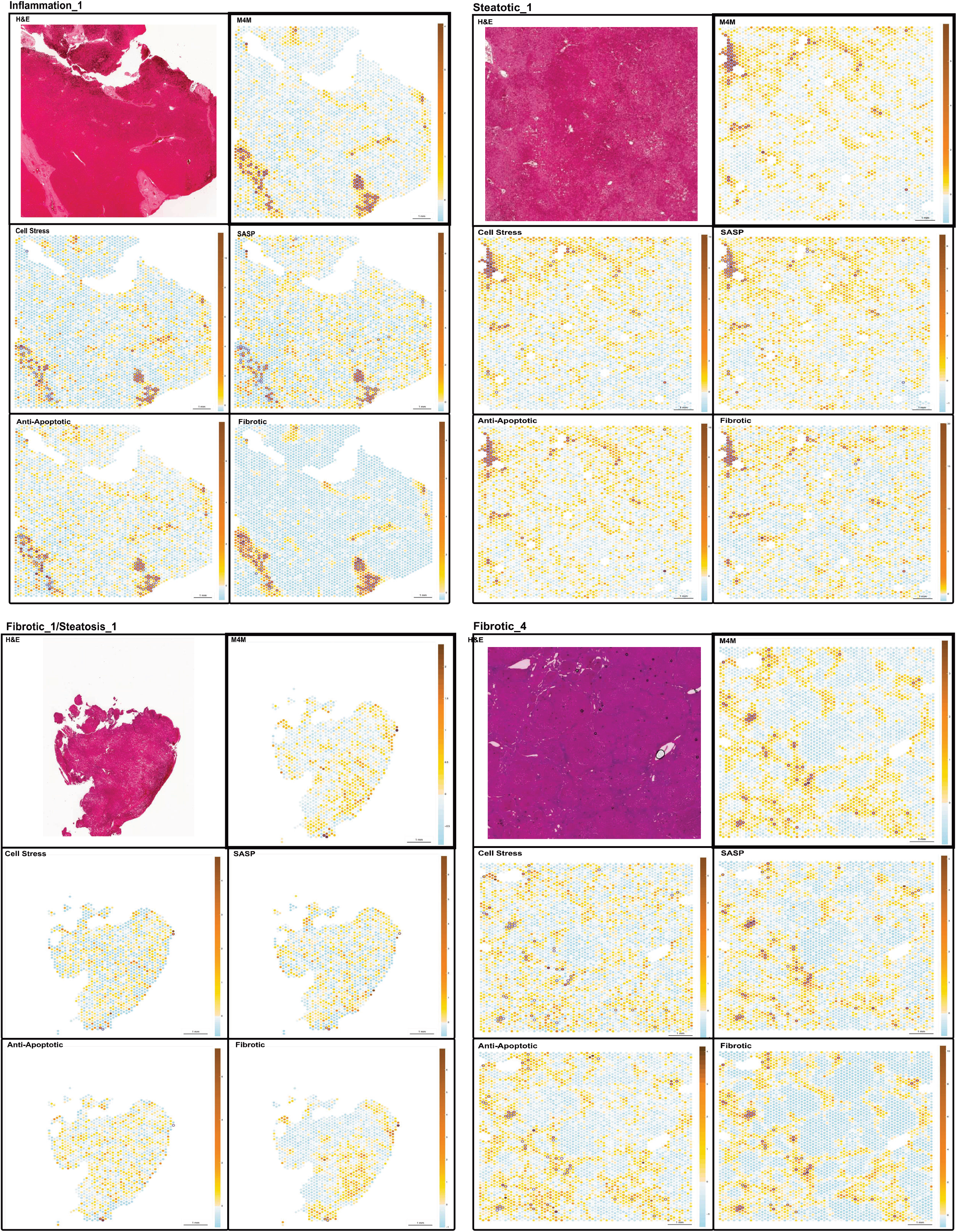
Images of Visium data identifying SenSpots in diseased liver samples. H&E (hematoxylin and eosin) stain of each tissue slide used for Visium, spatial heatmaps for M4M and the mean z-score for each senescence category: cell stress, SASP, anti-apoptotic, and fibrotic. SenSpots identified by M4M are highlighted with purple circles.

**Extended Data Figure 3.**
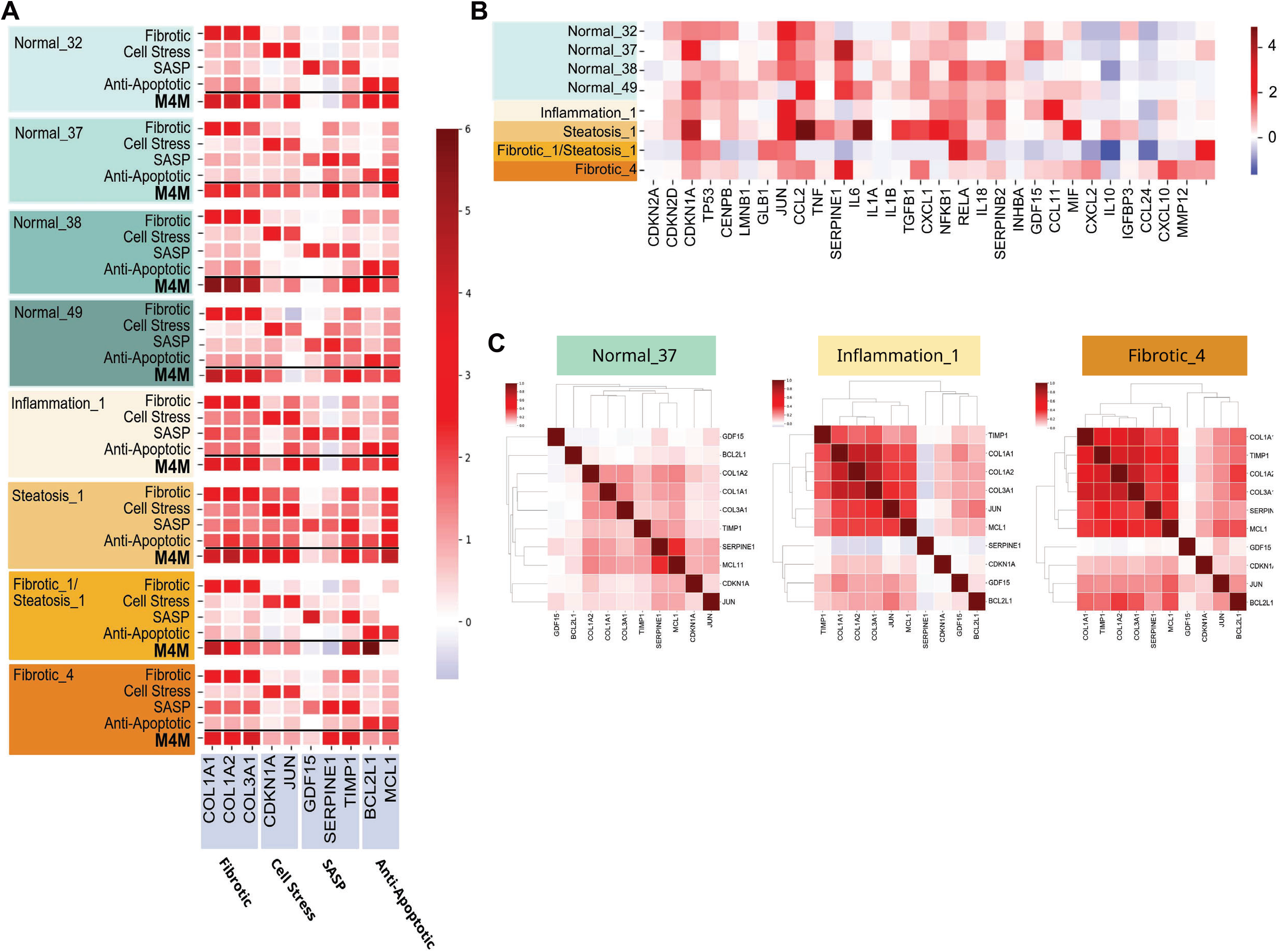
SenSpot Validation. **A.** Expression heat map for each of the 10 SenSpot-defining genes within alternative methods of niche identification. For each sample, alternative niches were identified using only enrichment of the single senescence hallmark category listed (fibrotic, cell stress, SASP, or anti-apoptotic) for each row. For comparison, integrated four-category identification of SenSpots by the mean of 4 means (M4M) method is shown for each sample under the black bar **B.** Expression of senescence-associated genes averaged across the SenSpots for each liver sample. **C.** The correlation of the 10 SenSpot-defining genes by stage of liver disease.

**Extended Data Figure 4.**
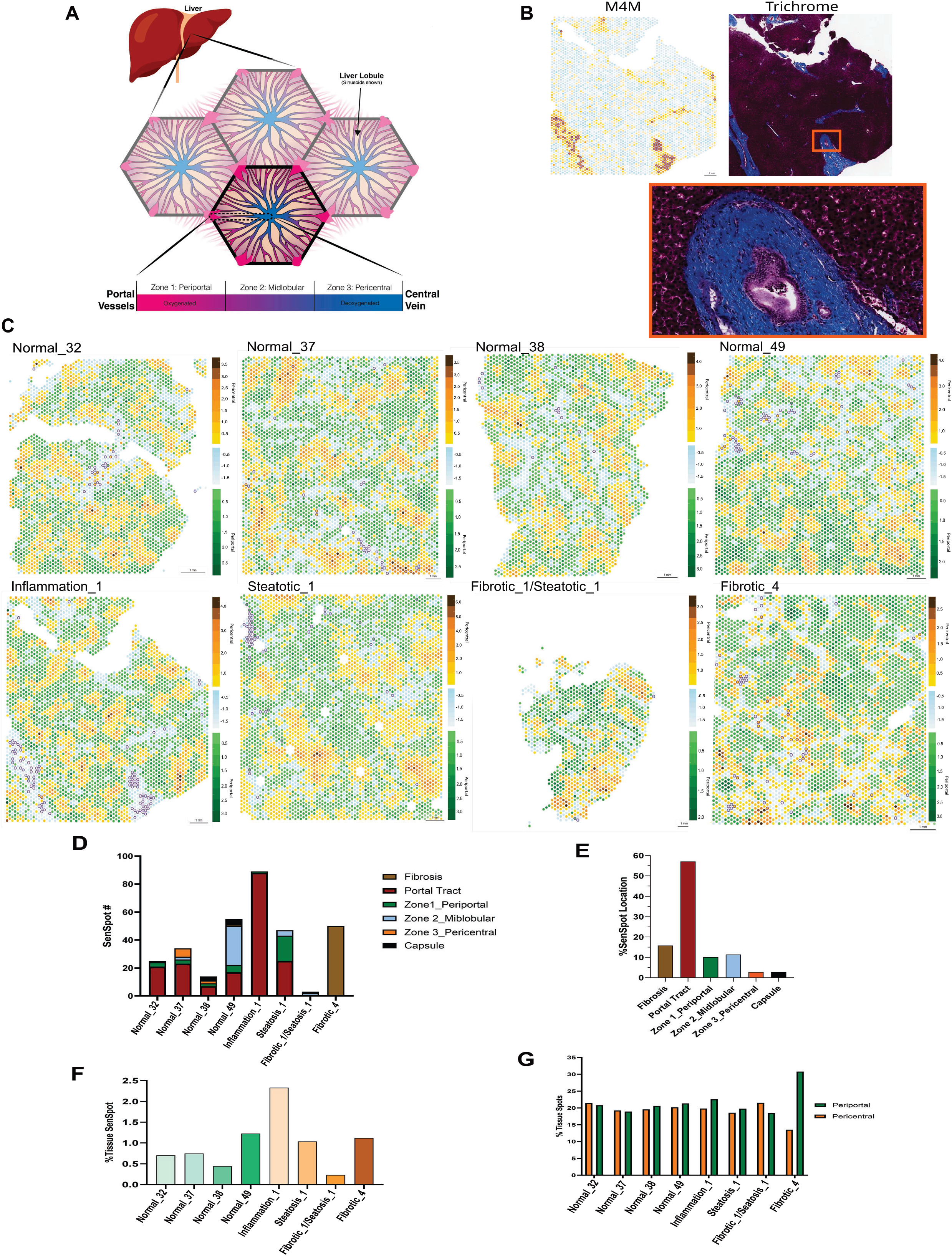
Localization of SenSpots. **A.** Schematic diagram of liver zonation **B.** Heatmap illustrating the SenSpots (purple circles) in the Inflammation_1 sample and a serial section with Trichrome staining to detect fibrosis (blue) 20X magnification enlarged to 200X. **C.** Spatial heatmaps of all 8 samples, indicating the pericentral region (yellow-brown) defined by *GLUL, CYP2E,* and *CYP27A1* expression, periportal region (green) defined by *ASL, ALB,* and *ASS1* expression, and the region with decreased expression of all 6 genes (blue). **D.** The anatomical location of SenSpots in liver mapped by a pathologist for each tissue sample. **E.** The percent of SenSpots identified in each liver anatomical region **F.** The percent of total Visium spots identified as a SenSpot for each tissue sample **G.** The percent of Visium spots identified as pericentral (orange) or periportal (green) for each tissue sample.

**Extended Data Figure 5.**
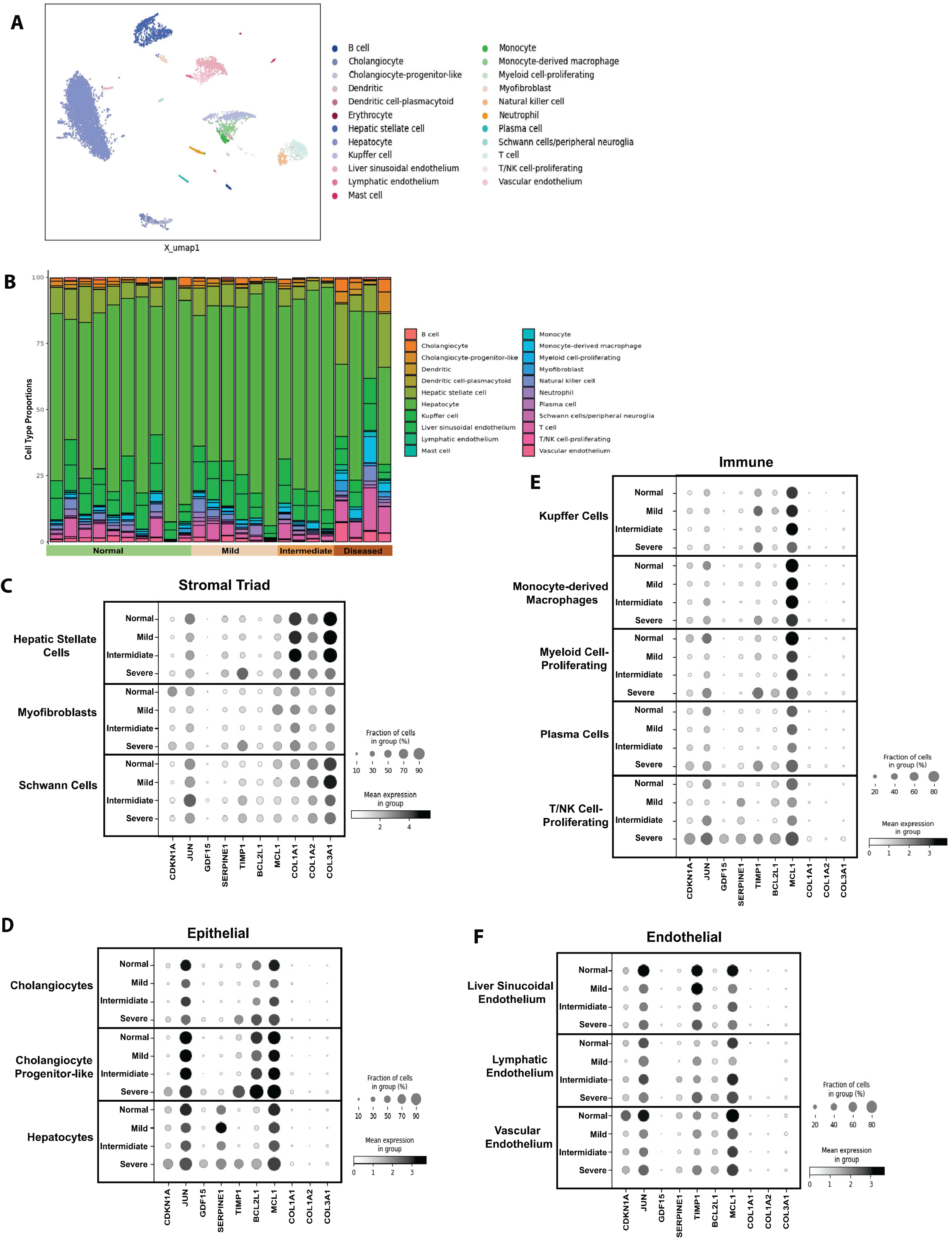
Single-cell RNAseq analysis of cell types with disease progression. **A.** UMAP representation of the cell-types identified in the scRNAseq data set. **B.** Cell-type composition (%) in each liver sample. **C-F.** Dot plots indicating the expression of the 10 M4M genes with increasing pathology from normal to severe in the indicated cell types in the scRNAseq data.

**Extended Data Figure 6:**
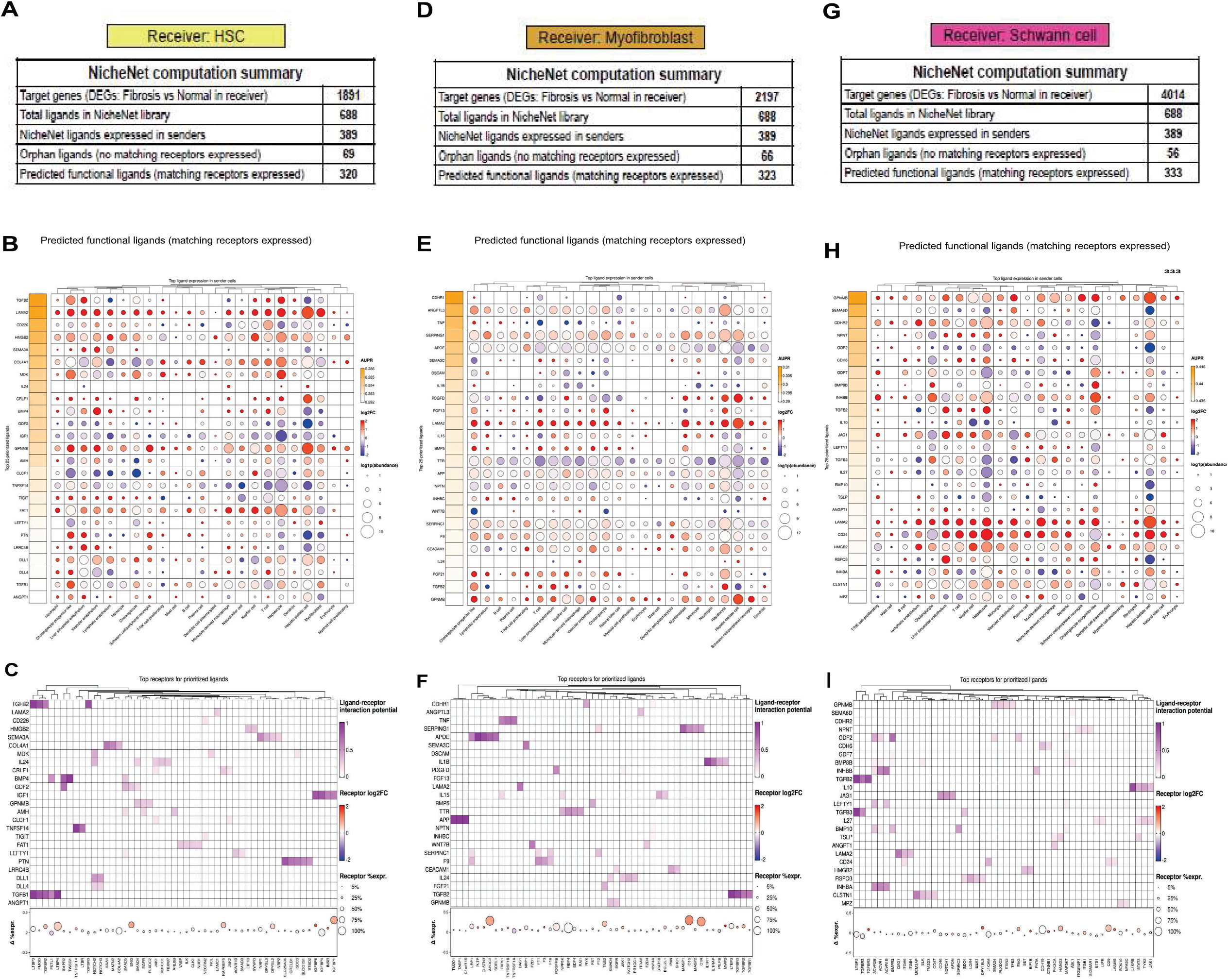
Expanded NicheNet ligand–receptor interactions across stromal triad receiver cell populations in fibrotic vs normal livers. **A-C.** NicheNet analysis of hepatic stellate cells (HSCs) as receiver cells. **A.** Summary of the analysis. **B.** Dot plot showing expression of ligands across potential sender cell-types. The dot size represents the ligand expression and the color indicates log2-fold-change in fibrotic vs normal liver. The ligands are ranked by area under the precision-recall curve (AUPR). **C.** Representation of the predicted interactions between ligands and the receptors expressed by HSCs. Purple represents the predicted ligand-receptor interaction potential. Orange circles represents the receptor log2 fold-change in fibrotic vs normal liver and the percentage of receiver cells expressing each receptor. **D-F.** Same as A-C but with myofibroblasts as receiver cells. **G-I.** Same as A-C but with Schwann cells as receiver cells.

**Extended Data Figure 7:**
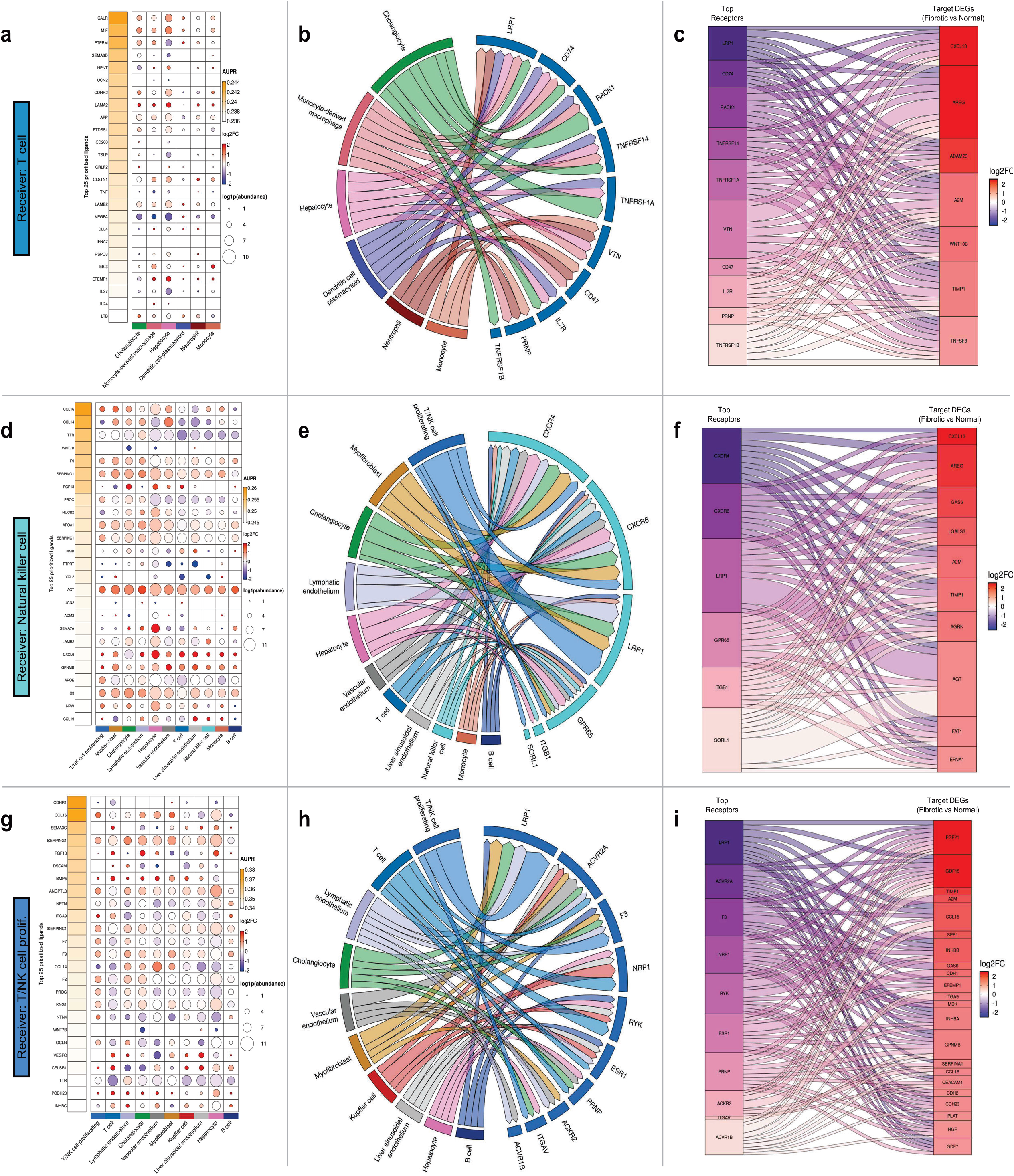
NicheNet modeling reveals regulation of T/NK cells in fibrotic livers. **A-C.** Schematic of the NicheNet analysis workflow used to interrogate cell-cell communication in fibrotic vs normal liver samples with T cells as receiver cells. **A.** Dot plot showing ligands by potential sender cell populations. The dot size represents ligand expression and the color indicates the log2 fold-change within the sender cell type in fibrotic vs normal livers. The ligands are ranked by area under the precision-recall curve (AUPR). **B.** Circos plot showing predicted interactions between sender cell populations and receiver cells with upregulated expression of potential receptors. The width of the connecting ribbons correlate to the weighted contribution of each sender cell to individual ligand-receptor interaction. **C.** Sankey plot linking T cell receptors to predicted downstream differentially expressed genes within receiver cells, colored by log2-fold-change in fibrotic vs normal livers. **D-F.** The same as A-C but for NK cells as potential receiver cells. **G-I.** The same as A-C but with proliferating T and NK cells as potential receiver cells.

**Extended Data Figure 8:**
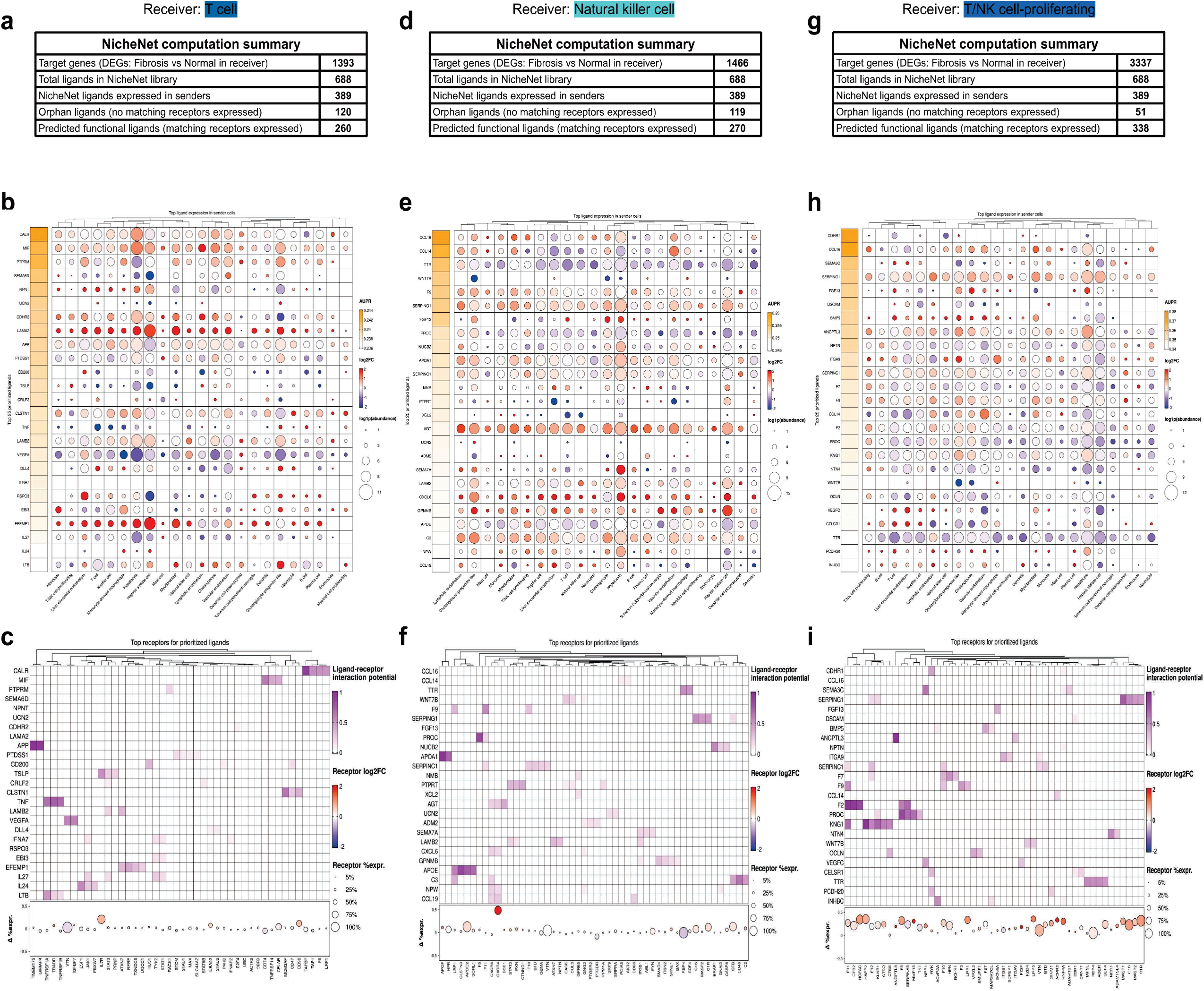
Expanded NicheNet ligand–receptor interactions across T and NK cell receiver populations in fibrotic vs normal livers. **A-C.** NicheNet analysis of T cells as receivers. **A.** Summary of analysis. **B.** Dot plot showing expression of ligands across potential sender cell-types. The dot size represents the ligand expression and the color indicates log2-fold change in fibrotic vs normal liver. The ligands are ranked by area under the precision-recall curve (AUPR). **C.** Representation of the predicted interactions between ligands and the receptors expressed by T cells. Purple represents the predicted ligand-receptor interaction potential. Orange circles represents the receptor log2-fold-change in fibrotic vs normal liver and the percentage of receiver cells expressing each receptor. **D-F.** Same as A-C but with NK cells as receivers. **G-I.** Same as A-C but with proliferating T and NK cells as receiver cells.

**Extended Data Figure 9.**
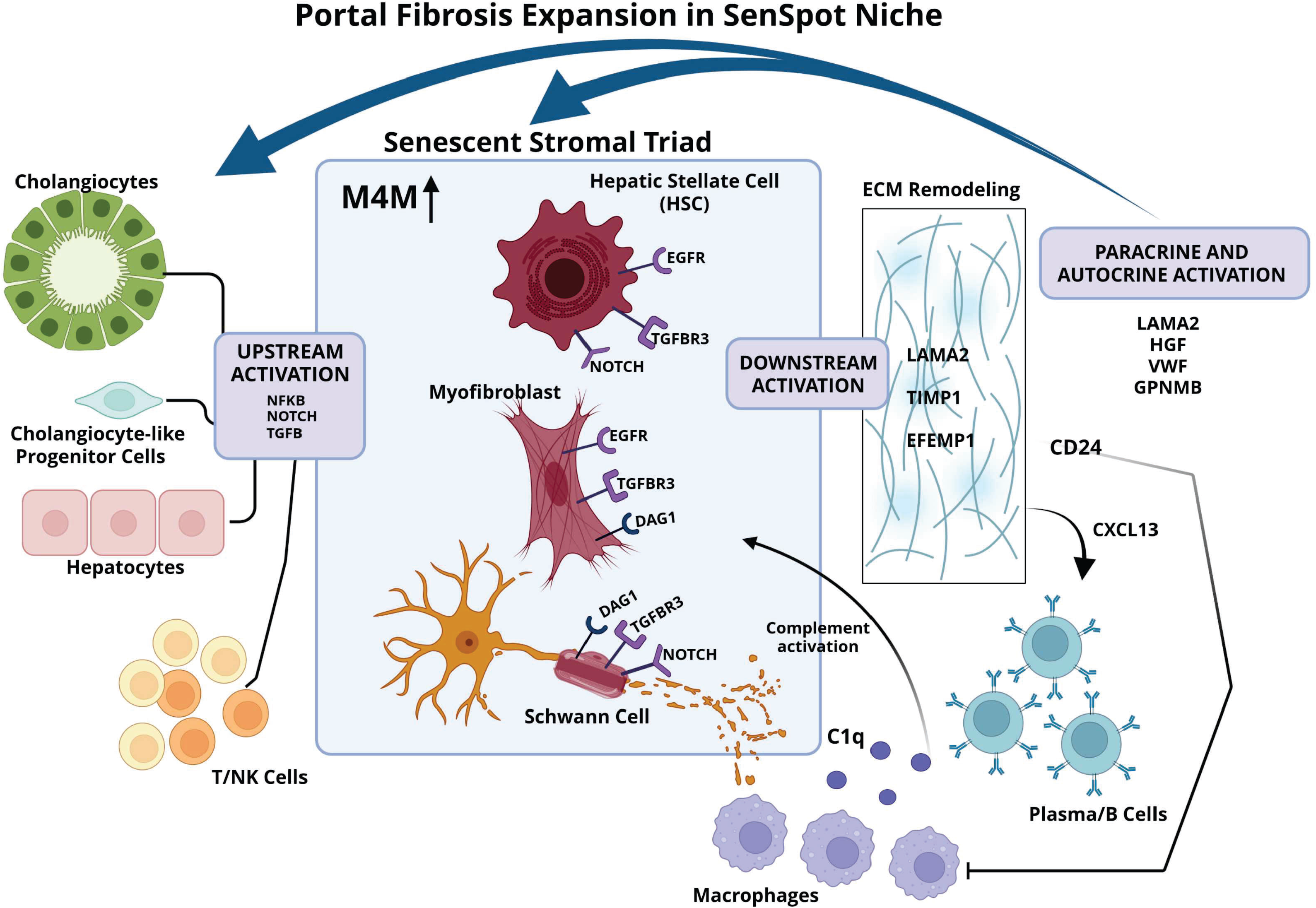
Portal fibrosis expansion in the SenSpot Niche. Upstream activation includes ligands from NF-κB, NOTCH, and TGF-β signaling pathways contributed by cholangiocytes, cholangiocyte-like progenitor cells, hepatocytes, and T or NK cells to the senescent stromal triad, which includes hepatic stellate cells, myofibroblasts, and Schwann cells. The stromal triad had upregulation of the receptors TGFβ3, EGF-R, NOTCH, and DAG1 leading to downstream activation of ECM remodeling genes (*LAMA2*, *TIMP1*, and *EFEMP1*) creating the basement membrane and foundation for fibrosis. Paracrine and autocrine activation occurring through LAMA2, HGF, VWF, and GPNMB increased with fibrotic disease. *CD24* expression from stromal cells inhibits macrophage phagocytosis and *CXCL13* recruits plasma/B cells to the fibrotic niche. Complement activation from macrophage expression of C1q and plasma/B cells immunoglobulin expression reinforces stromal cell senescence.

