## Supplemental Figures 1-3 and Supplemental Tables 1-4 for "Mapping Senescence Landscapes in Metabolic Liver Disease Progression"

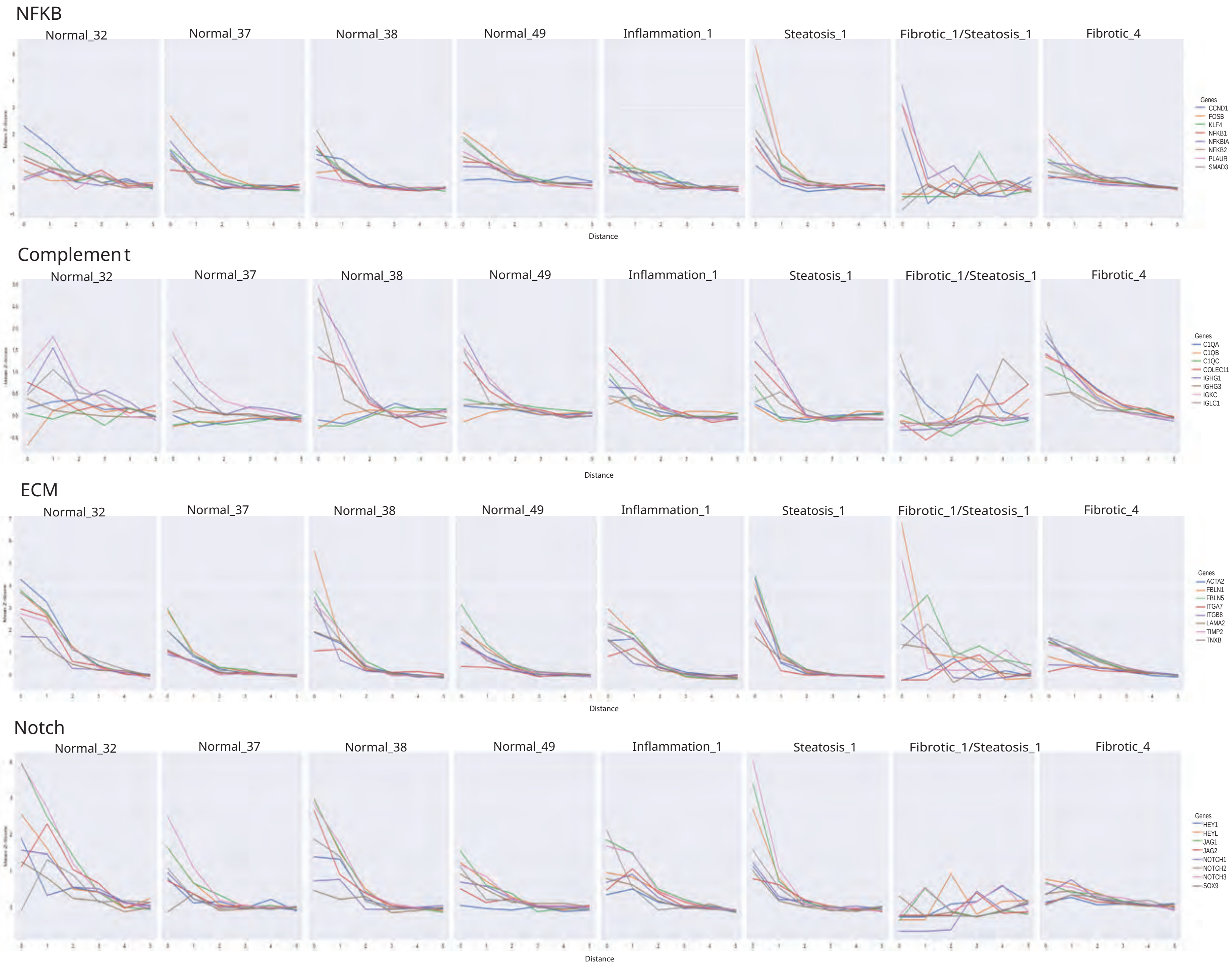
NFKB
Normal_32
Normal_37
Normal_38
Normal_49
Inflammation_1
Steatosis_1
Fibrotic_1/Steatosis_1
Fibrotic_4
Genes
CCND1
FOSB
KLF4
NFKB1
NFKBIA
NFKB2
PLAUR
SMAD3
Distance
Complement
Normal_32
Normal_37
Normal_38
Normal_49
Inflammation_1
Steatosis_1
Fibrotic_1/Steatosis_1
Fibrotic_4
Genes
C1QA
C1QB
C1QC
COLEC11
IGHG1
IGHG3
IGKC
IGLC1
Distance
ECM
Normal_32
Normal_37
Normal_38
Normal_49
Inflammation_1
Steatosis_1
Fibrotic_1/Steatosis_1
Fibrotic_4
Genes
ACTA2
FBLN1
FBLN5
ITGA7
ITGB8
LAMA2
TIMP2
TNXB
Distance
Notch
Normal_32
Normal_37
Normal_38
Normal_49
Inflammation_1
Steatosis_1
Fibrotic_1/Steatosis_1
Fibrotic_4
Genes
HEY1
HEYL
JAG1
JAG2
NOTCH1
NOTCH2
NOTCH3
SOX9
Distance

**Supplemental Figure 1.** Spatial Signal Expansion (SSE) analysis used to investigate the impact of SenSpots on their tissue microenvironment. SSE position 0 is the SenSpot and each successive Visium spot distance away from the SenSpot are distances 1-5. SSE graphs of SenSpot enriched pathways, NF-κB, complement, Notch signaling, and Extracellular Matrix (ECM) for all 8 liver samples.

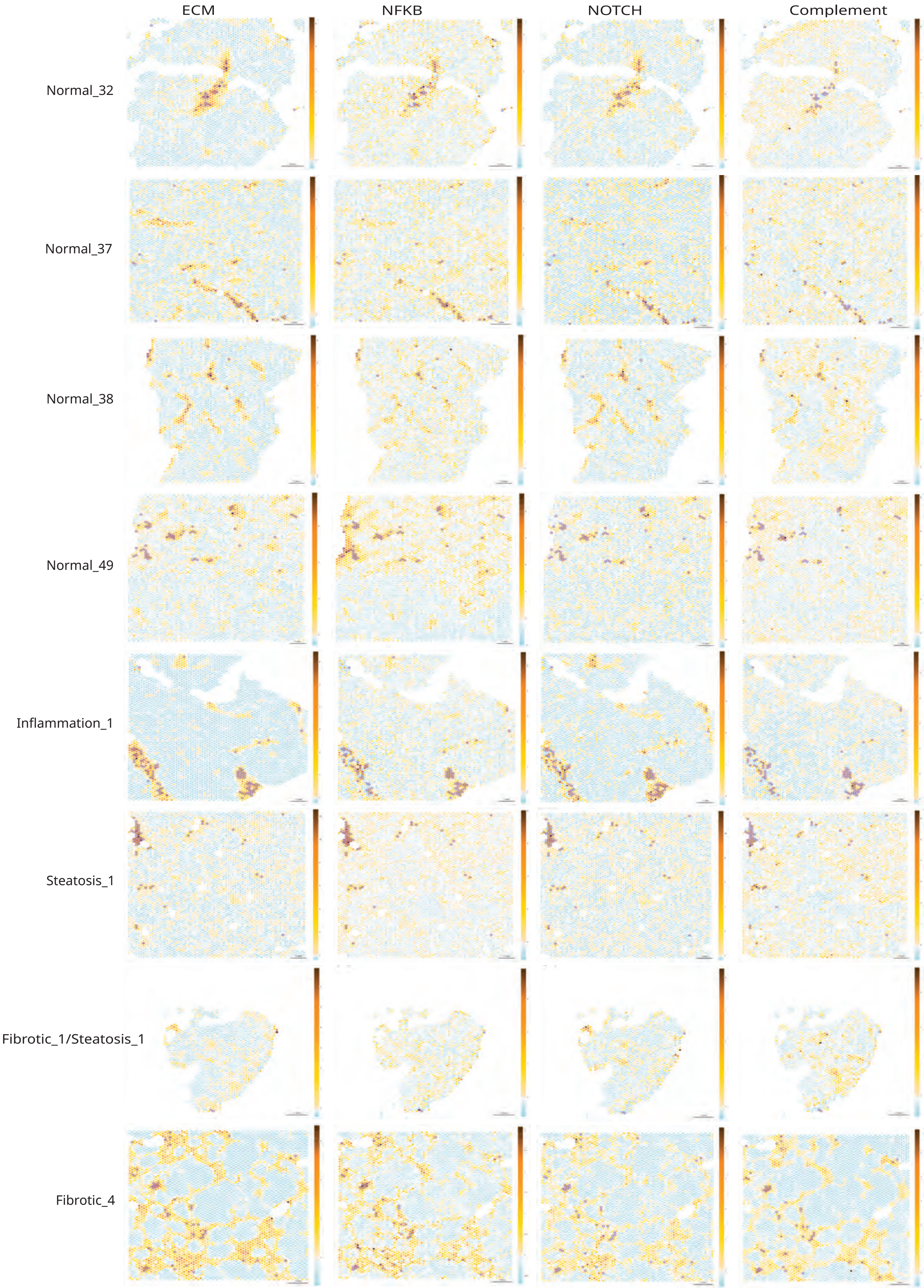

ECM
NFKB
NOTCH
Complement
Normal_32
Normal_37
Normal_38
Normal_49
Inflammation_1
Steatosis_1
Fibrotic_1/Steatosis_1
Fibrotic_4

**Supplemental Figure 2.** Spatial Signal Expansion (SSE) heatmaps of SenSpot enriched pathways, NF-κB, complement, Notch signaling, and Extracellular Matrix (ECM) for all liver samples labeled on left and pathway across the top. SenSpots are circled in purple.

SpaceRanger Analysis on Liver Visium CytAssist Data

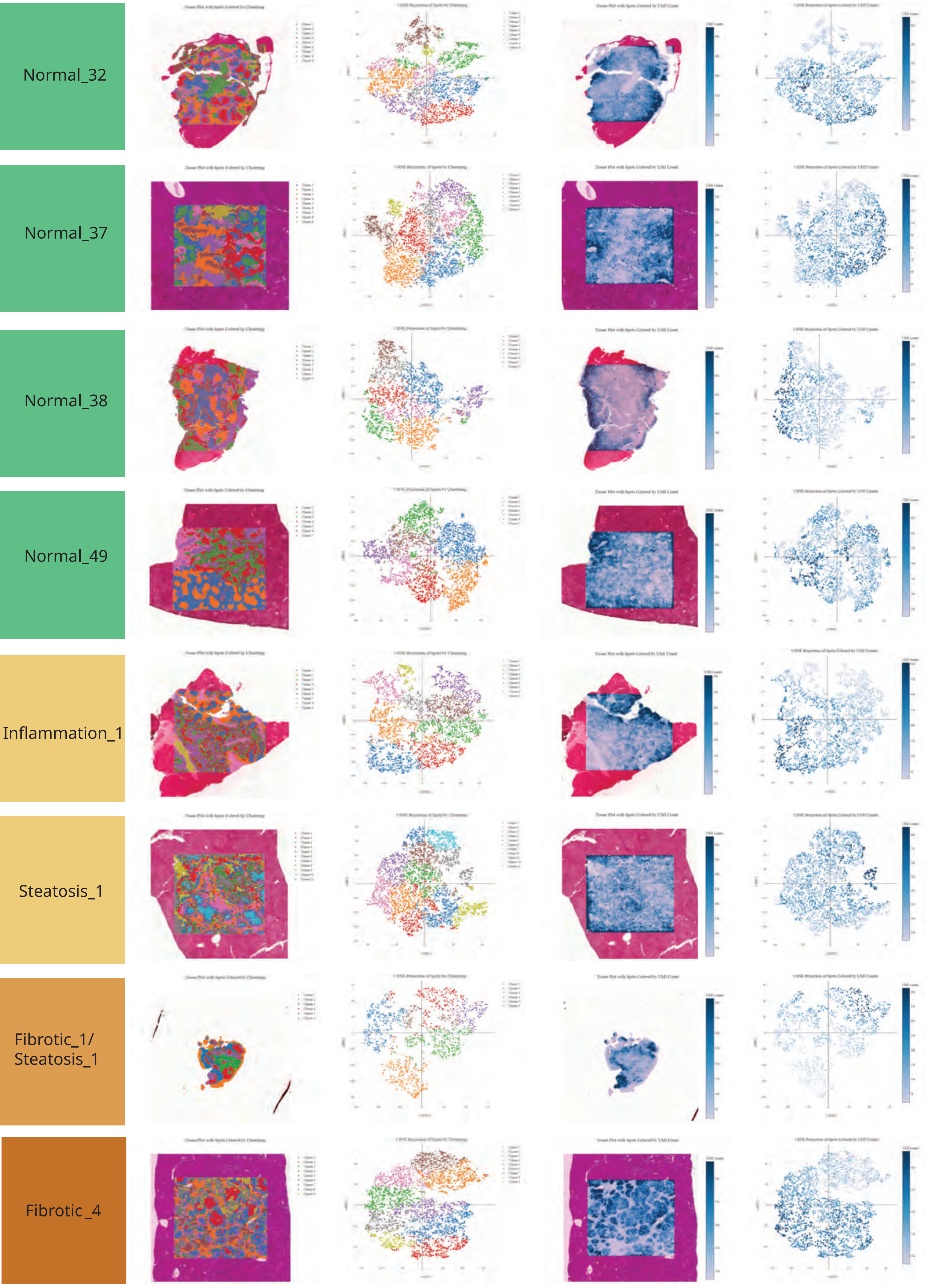

**Supplemental Figure 3.** 10x Genomics Space Ranger image outputs for each liver sample. **A.** Tissue plot with spots colored by clustering. **B.** tSNE projection of spots by clustering. **C.** Tissue plot with spots colored by UMI (unique molecular identifier) count. **D.** tSNE projection of spots colored by UMI counts.

**Supplemental Table 1.** 10x Genomics Visium CytAssist Space Ranger pipeline quality metric outputs

| Sample | Space Ranger Sample ID | Sample ID | Number of Spots Under Tissue | Number of Reads | Mean Reads per Spot | Mean Reads Under Tissue per Spot | Fraction of Spots Under Tissue | Valid Barcodes |
|---|---|---|---|---|---|---|---|---|
| Normal_32 | spaceranger_V42L29-278_D1 | BLS001684-1 | 4084 | 355426700 | 87029.0646 | 82948.30534 | 0.81810897 | 0.98793449 |
| Normal_37 | spaceranger_V42L06-053_D1 | T07-1450-11 | 4989 | 258971209 | 51908.4404 | 51163.93846 | 0.99939904 | 0.98675223 |
| Normal_38 | spaceranger_V42D20-289_D1 | BLS001648-0014 | 3436 | 445977013 | 129795.405 | 125445.459 | 0.68830128 | 0.99202687 |
| Normal_49 | spaceranger_V42L29-055_A1 | T10-0349-5 | 4907 | 653910656 | 133260.782 | 130357.0187 | 0.98297276 | 0.98925983 |
| Inflammation_1 | spaceranger_V42D20-289_A1 | BLS001460-0014 | 4199 | 505997686 | 120504.331 | 116673.1026 | 0.84114583 | 0.99215265 |
| Steatosis_1 | spaceranger_V42L29-055_D1 | T15-1102-14 | 4962 | 731318063 | 147383.729 | 145082.2707 | 0.99399038 | 0.98957684 |
| Fibrotic_1/ Steatosis_1 | spaceranger_V42L29-278_A1 | BLS001656-14 | 1593 | 147214417 | 92413.3189 | 81065.65599 | 0.31911058 | 0.98976502 |
| Fibrotic_4 | spaceranger_V42L06-053_A1 | T01-0006-19 | 4959 | 212998224 | 42951.85 | 42325.63319 | 0.99338942 | 0.98746647 |

| Sample | Valid UMIs | Sequencing Saturation | Q30 Bases in Barcode | Q30 Bases in Probe Read | Q30 Bases in UMI | Reads Mapped to Probe Set | Reads Mapped Confidently to Probe Set | Fraction Reads in Spots Under Tissue |
|---|---|---|---|---|---|---|---|---|
| Normal_32 | 0.992698931 | 0.641326203 | 0.957151751 | 0.95647721 | 0.96053506 | 0.98820535 | 0.981108046 | 0.965013513 |
| Normal_37 | 0.999850964 | 0.714420904 | 0.947837939 | 0.92550612 | 0.9489451 | 0.9729733 | 0.562459598 | 0.998200986 |
| Normal_38 | 0.999931384 | 0.739716256 | 0.967335466 | 0.94913142 | 0.96397017 | 0.98921545 | 0.985790431 | 0.97422809 |
| Normal_49 | 0.99987872 | 0.779680016 | 0.95467804 | 0.93971287 | 0.94848894 | 0.98508392 | 0.965466272 | 0.988700406 |
| Inflammation_1 | 0.999943464 | 0.851830541 | 0.967682978 | 0.95779587 | 0.96413429 | 0.99073041 | 0.985648322 | 0.97586406 |
| Steatosis_1 | 0.99984478 | 0.72306074 | 0.9545052 | 0.94101589 | 0.94788468 | 0.98635541 | 0.974504824 | 0.994867972 |
| Fibrotic_1/ Steatosis_1 | 0.992671913 | 0.785134118 | 0.96054058 | 0.95998187 | 0.96341002 | 0.98683094 | 0.898030816 | 0.892325679 |
| Fibrotic_4 | 0.999849111 | 0.516143402 | 0.948737182 | 0.92471482 | 0.94962852 | 0.9815863 | 0.97745877 | 0.997928102 |

| Sample | Median Genes per Spot | Median UMI Counts per Spot | Genes Detected | Reads Mapped Confidently to the Filtered Probe Set | Number of Genes | Number of Genes >= 10 UMIs | 10x Protocol CG000520 | 10x Protocol CG000495 |
|---|---|---|---|---|---|---|---|---|
| Normal_32 | 5355 | 24273.5 | 18055 | 0.956484594 | 18085 | 18016 | Revision B | Revision C |
| Normal_37 | 2633 | 7421 | 18054 | 0.551516593 | 18085 | 18000 | Revision B | Revision A |
| Normal_38 | 5692.5 | 24723.5 | 18057 | 0.95935373 | 18085 | 18007 | Revision B | Revision D |
| Normal_49 | 5113 | 24449 | 18063 | 0.949886896 | 18085 | 18030 | Revision B | Revision D |
| Inflammation_1 | 3779 | 13881 | 18044 | 0.963208749 | 18085 | 17996 | Revision B | Revision D |
| Steatosis_1 | 5730 | 34879.5 | 18056 | 0.957893181 | 18085 | 18022 | Revision B | Revision D |
| Fibrotic_1/ Steatosis_1 | 4227 | 11817 | 18050 | 0.879182302 | 18085 | 17985 | Revision B | Revision C |
| Fibrotic_4 | 4869 | 19252 | 18057 | 0.960629198 | 18085 | 18022 | Revision B | Revision A |

**Supplemental Table 2** Selection of senescence-associated genes used to identify SenSpots for each senescent cell feature category. Selected genes are highlighted in color: cellular stress (grey), senescence-associated secretory phenotype (SASP; orange), Anti-apoptotic (green), Fibrotic (teal)

| | Gene | Protein Name | median diseased | variance diseased | %nonzero spots diseased | median healthy | variance healthy | %nonzero spots healthy | disease-healthy |
|---|---|---|---|---|---|---|---|---|---|
| | CCNA2 | Cyclin A2 | 0 | 0.0786 | 0.06684 | 0 | 0.0565 | 0.0502 | 0.02213 |
| | CCNB2 | Cyclin B2 | 0 | 0.06401 | 0.0593 | 0 | 0.051 | 0.0441 | 0.01302 |
| | CDKN1A | Cyclin Dependent Kinase Inhibitor 1A | 2 | 11.21552 | **0.83411** | 2 | 4.9833 | **0.8454** | 6.23218 |
| | CDKN2A | Cyclin Dependent Kinase Inhibitor 2A | 0 | 0.03019 | 0.0269 | 0 | 0.0304 | 0.0282 | -0.0002 |
| | CDKN2D | Cyclin Dependent Kinase Inhibitor 2D | 0 | 0.1315 | 0.11351 | 0 | 0.1257 | 0.1086 | 0.00581 |
| | CENPB | Centromere Protein B | 0 | 0.71287 | 0.39209 | 0 | 0.9668 | 0.495 | -0.25398 |
| **Cellular Stress** | GLB1 | Beta Galactosidase 1 | 1 | 1.09511 | **0.51844** | 1 | 2.2673 | **0.7235** | -1.17214 |
| | JUN | Jun Proto-Oncogene | 19 | 175.63925 | **0.99878** | 5 | 26.9679 | **0.9851** | 148.67131 |
| | LMNB1 | Lamin-B1 | 0 | 0.37739 | 0.25576 | 0 | 0.5036 | 0.3444 | -0.1262 |
| | MKI67 | Marker of Proliferation KI-67 | 0 | 0.14639 | 0.10964 | 0 | 0.0765 | 0.0558 | 0.06989 |
| | STMN1 | Stathmin 1 | 0 | 0.04543 | 0.04443 | 0 | 0.0456 | 0.0397 | -0.00017 |
| | TP53 | Tumor Protein P53 | 0 | 0.78726 | 0.43122 | 1 | 1.3321 | **0.6028** | -0.54483 |
| | TP53BP1 | Tumor Protein p53 Binding Protein 1 | 0 | 0.19756 | 0.16038 | 0 | 0.2256 | 0.1906 | -0.028 |
| | CCL2 | C-C Motif Chemokine Ligand 2 | 0 | 7.42405 | 0.41451 | 1 | 6.0131 | **0.6614** | 1.41095 |
| | CCL11 | C-C Motif Chemokine Ligand 11 | 0 | 0.0329 | 0.02914 | 0 | 0.0702 | 0.0385 | -0.0373 |
| | CCL24 | C-C Motif Chemokine Ligand 24 | 0 | 0.08028 | 0.06908 | 0 | 0.1421 | 0.0985 | -0.0618 |
| | CXCL1 | C-X-C Motif Chemokine Ligand 1 | 0 | 0.1127 | 0.07112 | 0 | 0.1773 | 0.1157 | -0.06459 |
| | CXCL10 | C-X-C Motif Chemokine Ligand 10 | 0 | 6.88128 | 0.26411 | 0 | 13.8835 | 0.1747 | -7.00224 |
| | CXCL2 | C-X-C Motif Chemokine Ligand 2 | 6 | 32.32414 | **0.98655** | 15 | 81.652 | **0.9988** | -49.32787 |
| **SASP** | GDF15 | Growth Differentiation Factor 15 | 3 | 24.62516 | **0.84206** | 0 | 0.888 | 0.394 | 23.73717 |
| | HMGB1 | High Mobility Group Box 1 | | | 0 | | | 0 | 0 |
| | IGFBP3 | Insulin Like Growth Factor Binding Proten 3 | 10 | 85.53844 | **0.99389** | 10 | 98.258 | **0.9948** | -12.71953 |
| | IL10 | Interleukin 10 | 0 | 0.09293 | 0.07438 | 0 | 0.0488 | 0.0455 | 0.04417 |
| | IL18 | Interleukin 18 | 0 | 0.43209 | 0.28327 | 0 | 0.1926 | 0.162 | 0.23951 |
| | IL1A | Interleukin 1 alpha | 0 | 0.03505 | 0.03444 | 0 | 0.0362 | 0.0351 | -0.00115 |
| | IL1B | Interleukin 1 beta | 0 | 0.13428 | 0.0966 | 0 | 0.4187 | 0.2346 | -0.28444 |
| | IL6 | Interleukin 6 | 0 | 0.3009 | 0.07826 | 0 | 0.4005 | 0.0943 | -0.09962 |
| | INHBA | Activin A/Inhibin Subunit Beta A | 0 | 0.99608 | 0.4677 | 1 | 1.7959 | **0.663** | -0.79986 |

| | | | | | | | | | |
|---|---|---|---|---|---|---|---|---|---|
| | MIF | Macrophage Migration Inhibitory Factor | 1 | 3.67167 | **0.75158** | 3 | 12.2695 | **0.949** | -8.59784 |
| | MMP12 | Matrix Metallopeptidase 12 | 0 | 0.03786 | 0.03811 | 0 | 0.0509 | 0.0425 | -0.01306 |
| | NFKB1 | Nuclear Factor Kappa B Subunit 1 | 1 | 2.60816 | **0.68881** | 1 | 2.0071 | **0.6884** | 0.60106 |
| | RELA | RELA Pronto-Oncogene, NF-KB Subunit | 1 | 1.3399 | **0.56735** | 1 | 1.3017 | **0.5959** | 0.03815 |
| | SERPINB2 | Plasminogen Activator Inhibitor 2 | 0 | 0.03843 | 0.03098 | 0 | 0.0348 | 0.0318 | 0.00364 |
| | SERPINE1 | Plasminogen Activator Inhibitor 1 | 19 | 651.5202 | **0.99817** | 36 | 1016.3152 | **0.9994** | -364.79498 |
| | TGFB1 | Transforming Growth Factor Beta 1 | 0 | 0.36103 | 0.21031 | 0 | 0.1972 | 0.1397 | 0.16379 |
| | TIMP1 | TIMP Metallopeptidase Inhibitor 1 | 4 | 40.82175 | **0.95741** | 2 | 8.9918 | **0.8769** | 31.82998 |
| | TIMP2 | TIMP Metallopeptidase Inhibitor 2 | 2 | 5.54187 | **0.84858** | 1 | 2.7065 | **0.6727** | 2.83534 |
| | TNF | Tumor Necrosis Factor alpha | 0 | 0.08572 | 0.07622 | 0 | 0.0999 | 0.0836 | -0.01417 |
| **Anti-Apoptotic** | BCL2 | BCL2 Apoptosis Regulator | 0 | 0.48321 | 0.28143 | 0 | 0.2866 | 0.2082 | 0.19665 |
| | BCL2A1 | BCL2A1, BCL2L5, BFL1 | 0 | 0.26483 | 0.16405 | 0 | 0.4998 | 0.2989 | -0.23493 |
| | BCL2L1 | BCL2L1, BCLXL | 2 | 5.71013 | **0.85164** | 2 | 3.9165 | **0.839** | 1.79361 |
| | BCL2L10 | Boo, Diva | 0 | 0.2855 | 0.20522 | 0 | 1.2961 | 0.4992 | -1.01056 |
| | BCL2L2 | BCL2L2 | 0 | 0.63193 | 0.35419 | 0 | 0.6548 | 0.3962 | -0.02288 |
| | MCL1 | BCL2L3, MCL1L | 26 | 535.49268 | **0.99898** | 17 | 180.7291 | **0.9996** | 354.76355 |
| **Fibrotic** | COL1A1 | Collagen Type I Alpha 1 chain | 2 | 17.74072 | **0.82719** | 1 | 4.6771 | **0.6526** | 13.06358 |
| | COL1A2 | Collagen Type I Alpha 2 Chain | 1 | 7.41495 | **0.69003** | 1 | 2.428 | **0.5476** | 4.98698 |
| | COL3A1 | Collagen Type III Alpha 1 Chain | 7 | 72.3557 | **0.9841** | 4 | 21.3631 | **0.9708** | 50.9926 |
| | MMP1 | Matrix metallopeptidase 1 | 0 | 0.02508 | 0.02384 | 0 | 0.0307 | 0.0304 | -0.00561 |
| | MMP2 | Matrix metallopeptidase 2 | 0 | 1.06381 | 0.39128 | 0 | 0.5426 | 0.2233 | 0.52122 |
| | MMP9 | Matrix metallopeptidase 9 | 0 | 0.17504 | 0.0485 | 0 | 0.5424 | 0.3049 | -0.36738 |
| | PDGFA | Platelet Derived Growth Factor Subunit A | 0 | 0.49455 | 0.25596 | 0 | 0.1857 | 0.1306 | 0.30882 |
| | PDGFB | Platelet Derived Growth Factor Subunit A | 0 | 0.30549 | 0.21296 | 0 | 0.2234 | 0.152 | 0.08213 |

**Supplemental Table 3.** Pathology grading of the liver samples used for scRNAseq including five of the Visium CytAssist samples.

| Visium Sample | Specimen/ Subject ID | Patient Age | Steatosis, macro % | Steatosis score (0, 1, 2, 3) | Ballooning (0, 1, 2) | Inflammation (0, 1, 2, 3) | Fibrosis (0, 1a, 1b, 1c, 2, 3, 4) |
|---|---|---|---|---|---|---|---|
| | T14-0545-10 | 29 | 0 | 0 | 0 | 0 | 0 |
| | BLS001607 | 30 | 0 | 0 | 0 | 0 | 0 |
| Normal_38 | BLS001648 | 38 | 1 | 0 | 0 | 0 | 0 |
| Normal_37 | T07-1450-11 | 37 | 0 | 0 | 0 | 0 | 0 |
| | T99-1476-19 | 38 | 0 | 0 | 0 | 0 | 0 |
| | T09-2087-4 | 46 | 0 | 0 | 0 | 0 | 0 |
| Normal_49 | T10-0349-5 | 49 | 0 | 0 | 0 | 0 | 0 |
| | BLS001555 | 43 | 2 | 0 | 0 | 0 | 0 |
| | T14-0183-3 | 66 | 5 | 0 | 0 | 0 | 0 |
| | BLS001625 | 60 | 5 | 0 | 0 | 0 | 0 |
| | BLS001572 | 51 | 1 | 0 | 0 | 1 | 0 |
| Steatosis_1 | T15-1102-14 | 50 | 10 | 1 | 0 | 0 | 0 |
| | T14-0106-4 | 52 | 10 | 1 | 0 | 0 | 0 |
| | T18-0493-8 | 59 | 10 | 1 | 0 | 0 | 0 |
| | BLS001606 | 42 | 15 | 1 | 0 | 0 | 0 |
| | BLS001947 | 42 | 15 | 1 | 0 | 0 | 0 |
| | BLS001946 | 57 | 35 | 2 | 1 | 1 | 0 |
| | T18-0748-4 | 30 | 40 | 2 | 1 | 1 | 0 |
| | T17-1330-8 | 67 | 10 | 1 | 1 | 2 | 0 |
| | T19-0072-52 | 51 | 20 | 1 | 1 | 1 | 1a |
| | T02-0058-20 | 38 | 0 | 0 | 0 | 0 | 4 |
| Fibrotic_4 | T01-0006-19 | 50 | 0 | 0 | 0 | 1 | 4 |
| | T01-0512-20 | 43 | 0 | 0 | 0 | 0 | 3 |
| | T99-1433-20 | 55 | 5 | 1 | 1 | 1 | 4 |

**Supplemental Table 4.** Data Access through SenNet Consortium

### Visium Samples

| Pathology | Sample | Specimen/ Subject ID | Patient Age | Single-Cell Datasets (10x RNA Flex, Fix/Dis) | |
|---|---|---|---|---|---|
| | | | | SenNet ID | DOI URL |
| Normal | Normal_32 | BLS001684 | 32 | SNT952.XZZN.754 | https://doi.org/10.60586/SNT952.XZZN.754 |
| | Normal_37 | T07-1450-11 | 37 | SNT932.SKQQ.338 | https://doi.org/10.60586/SNT932.SKQQ.338 |
| | Normal_38 | BLS001648 | 38 | SNT497.JNCZ.552 | https://doi.org/10.60586/SNT497.JNCZ.552 |
| | Normal_49 | T10-0349-5 | 49 | SNT545.XPTS.687 | https://doi.org/10.60586/SNT545.XPTS.687 |
| Mild | Inflammation_1 | BLS001460 | 42 | SNT968.JQNX.889 | https://doi.org/10.60586/SNT968.JQNX.889 |
| | Steatosis_1 | T15-1102-14 | 50 | SNT379.NTWK.593 | https://doi.org/10.60586/SNT379.NTWK.593 |
| Intermediate | Fibrotic_1/ Steatosis_1 | BLS001656 | 42 | SNT697.TFFT.696 | https://doi.org/10.60586/SNT697.TFFT.696 |
| Diseased | Fibrotic_4 | T01-0006-19 | 50 | SNT433.JVGW.447 | https://doi.org/10.60586/SNT433.JVGW.447 |

### Single Cell RNAseq Samples

| Pathology | Visium Sample | Specimen/ Subject ID | Patient Age | Single-Cell Datasets (10x RNA Flex, Fix/Dis) | |
|---|---|---|---|---|---|
| | | | | SenNet ID | DOI URL |
| Normal | | T14-0545-10 | 29 | SNT568.LQMW.245 | https://doi.org/10.60586/SNT568.LQMW.245 |
| | | BLS001607 | 30 | SNT249.ZJSG.526 | https://doi.org/10.60586/SNT249.ZJSG.526 |
| | Normal_38 | BLS001648 | 38 | SNT743.NXQJ.774 | https://doi.org/10.60586/SNT743.NXQJ.774 |
| | Normal_37 | T07-1450-11 | 37 | SNT659.MFCT.299 | https://doi.org/10.60586/SNT659.MFCT.299 |
| | | T99-1476-19 | 38 | SNT928.KQBD.339 | https://doi.org/10.60586/SNT928.KQBD.339 |
| | | T09-2087-4 | 46 | SNT426.GBPP.347 | https://doi.org/10.60586/SNT426.GBPP.347 |
| | Normal_49 | T10-0349-5 | 49 | SNT298.TGFD.448 | https://doi.org/10.60586/SNT298.TGFD.448 |
| | | BLS001555 | 43 | SNT827.RSQX.548 | https://doi.org/10.60586/SNT827.RSQX.548 |
| | | T14-0183-3 | 66 | SNT625.XWKW.494 | https://doi.org/10.60586/SNT625.XWKW.494 |
| | | BLS001625 | 60 | SNT257.FNNC.672 | https://doi.org/10.60586/SNT257.FNNC.672 |
| Mild | | BLS001572 | 51 | SNT899.ZMQK.332 | https://doi.org/10.60586/SNT899.ZMQK.332 |
| | Steatosis_1 | T15-1102-14 | 50 | SNT296.FFXG.687 | https://doi.org/10.60586/SNT296.FFXG.687 |
| | | T14-0106-4 | 52 | SNT732.TJCR.969 | https://doi.org/10.60586/SNT732.TJCR.969 |
| | | T18-0493-8 | 59 | SNT943.QTNB.243 | https://doi.org/10.60586/SNT943.QTNB.243 |
| | | BLS001606 | 42 | SNT366.FRLM.998 | https://doi.org/10.60586/SNT366.FRLM.998 |
| | | BLS001947 | 42 | SNT573.SXDQ.942 | https://doi.org/10.60586/SNT573.SXDQ.942 |
| Intermediate | | BLS001946 | 57 | SNT587.VWSB.325 | https://doi.org/10.60586/SNT587.VWSB.325 |
| | | T18-0748-4 | 30 | SNT445.PQVM.734 | https://doi.org/10.60586/SNT445.PQVM.734 |
| | | T17-1330-8 | 67 | SNT232.ZWGG.323 | https://doi.org/10.60586/SNT232.ZWGG.323 |
| | | T19-0072-52 | 51 | SNT444.HPBV.724 | https://doi.org/10.60586/SNT444.HPBV.724 |
| Diseased | | T02-0058-20 | 38 | SNT634.FNPP.234 | https://doi.org/10.60586/SNT634.FNPP.234 |
| | Fibrotic_4 | T01-0006-19 | 50 | SNT329.ZLMR.757 | https://doi.org/10.60586/SNT329.ZLMR.757 |
| | | T01-0512-20 | 43 | SNT569.XDSQ.484 | https://doi.org/10.60586/SNT569.XDSQ.484 |
| | | T99-1433-20 | 55 | SNT782.THPT.548 | https://doi.org/10.60586/SNT782.THPT.548 |
